# A transient role of primary cilia in controlling direct versus indirect neurogenesis in the developing cerebral cortex

**DOI:** 10.1101/2020.04.28.065615

**Authors:** Kerstin Hasenpusch-Theil, Christine Laclef, Matt Colligan, Eamon Fitzgerald, Katherine Howe, Emily Carroll, Shaun R. Abrams, Jeremy F. Reiter, Sylvie Schneider-Maunoury, Thomas Theil

## Abstract

During the development of the cerebral cortex, neurons are generated directly from radial glial cells or indirectly via basal progenitors. The balance between these division modes determines the number and types of neurons formed in the cortex thereby affecting cortical functioning. Here, we investigate the role of primary cilia in this process. We show that a mutation in the ciliary gene *Inpp5e* leads to a transient increase in direct neurogenesis and subsequently to an overproduction of layer V neurons in newborn mice. Loss of *Inpp5e* also affects ciliary structure coinciding with increased Akt and mTOR signalling and reduced Gli3 repressor levels. Genetically re-storing Gli3 repressor rescues the decreased indirect neurogenesis in *Inpp5e* mutants. Overall, our analyses reveal how primary cilia determine neuronal subtype composition of the cortex by controlling direct vs indirect neurogenesis. These findings have implications for understanding cortical malformations in ciliopathies with *INPP5E* mutations.

## INTRODUCTION

Building a functional cerebral cortex which confers humans with their unique cognitive capabilities requires controlling the proliferation of neural progenitor cells and the timing and modes of neurogenic cell divisions. Varying the timing and modes of neurogenesis affects neuronal numbers and subtype composition of the cortex (Florio & Huttner, 2014). In the developing murine cortex, radial glial cells (RGCs) represent the major neural stem cell type. Residing in the ventricular zone, they express Pax6 and undergo interkinetic nuclear migration dividing at the ventricular surface (Götz, Stoykova, & Gruss, 1998; Warren et al., 1999). Initially, RGCs go through rounds of symmetric proliferative divisions to produce two RGCs increasing the progenitor pool but switch to asymmetric divisions at the beginning of cortical neurogenesis. RGCs generate neurons in two ways, either directly or indirectly via the production of basal progenitors (BPs) that settle in the subventricular zone (SVZ) and express the Tbr2 transcription factor (Englund et al., 2005). In the mouse, the majority of BPs divide once to produce two neurons whereas the remainders undergo one additional round of symmetric proliferative division before differentiating into two neurons (Haubensak, Attardo, Denk, & Huttner, 2004; Miyata et al., 2004; Noctor, Martinez-Cerdeno, Ivic, & Kriegstein, 2004). In this way, BPs increase neuron output per radial glial cell and have therefore been implicated in the evolutionary expansion of the mammalian cerebral cortex (Martinez-Cerdeno, Noctor, & Kriegstein, 2006). Thus, the balance between direct and indirect neurogenesis is an important factor in generating appropriate neuron numbers and types.

The mechanisms that fine tune this balance and thereby adjust the numbers and types of neurons produced in the cortex have only recently been investigated. Mitotic spindle orientation (Postiglione et al., 2011) and endoplasmic reticulum (ER) stress (Gladwyn-Ng et al., 2018; Laguesse et al., 2015) are contributing factors to control the generation of basal progenitors. In addition, levels of Slit/Robo and Notch/Delta signaling were shown to be evolutionarily conserved factors that determine the predominant mode of neurogenesis (Cardenas et al., 2018). Moreover, feedback signals from postmitotic neurons control the fate of radial glial daughter cells involving the release of Neurotrophin-3 and Fgf9 (Parthasarathy, Srivatsa, Nityanandam, & Tarabykin, 2014; Seuntjens et al., 2009) as well as the activation of a Notch-dependent signaling pathway (W. Wang et al., 2016). These studies highlight the importance of cell-cell signaling in controlling the cell lineage of cortical progenitors (Silva, Peyre, & Nguyen, 2019) and emphasize the necessity of studying the cellular mechanisms by which these signals control the decision by RGCs to undergo direct or indirect neurogenesis.

Given the importance of cell-cell signaling, it is likely that the primary cilium, a signaling hub in embryogenesis in general and in neural development in particular (Valente, Rosti, Gibbs, & Gleeson, 2014), plays key roles in determining the balance between direct versus indirect neurogenesis. The cilium is a subcellular protrusion that predominately emanates from the apical surface of radial glial cells projecting into the ventricular lumen. The phenotypes of several mouse lines mutant for ciliary genes underline the importance of the primary cilium in forebrain development but these mutants often suffer from severe patterning defects (Ashique et al., 2009; Besse et al., 2011; Willaredt et al., 2008) which make studying ciliary roles in determining the lineage of cortical progenitors difficult. To address such functions, we investigated corticogenesis in a mouse mutant for the ciliary gene *Inpp5e*.

*INPP5E* is mutated in Joubert syndrome (JS) (Bielas et al., 2009; Jacoby et al., 2009), a ciliopathy characterized by cerebellar defects in which a subset of patients also shows malformations of the cerebral cortex including heterotopias, polymicrogyria and agenesis of the corpus callosum (Valente et al., 2014). *Inpp5e* encodes Inositol polyphosphate 5 phosphatase E, an enzyme that is localized in the ciliary membrane and that hydrolyses the phosphatidylinositol polyphosphates PI(4,5)P_2_ and PI(3,4,5)P_3_ (Bielas et al., 2009; Jacoby et al., 2009). In this way, it controls the inositol phosphate composition of the ciliary membrane and thereby regulates the activity of several signaling pathways and cilia stability (Bielas et al., 2009; Chavez et al., 2015; Garcia-Gonzalo et al., 2015; Jacoby et al., 2009; Plotnikova et al., 2015). In contrast to *Inpp5e*’s extensively studied biochemical and cellular roles, little is known how these diverse functions are employed at the tissue level to control RGC lineage.

Here, we show that loss of *Inpp5e* function results in an increase in neuron formation at the expense of basal progenitor production in the E12.5 cortex and in an overproduction of Ctip2+ layer V neurons in newborn mutants. Moreover, RGC cilia show unusual membranous structures and/or abnormal numbers of microtubule doublets affecting the signaling capabilities of the cilium. The levels of Gli3 repressor (Gli3R), a critical regulator of cortical stem cell development (Hasenpusch-Theil et al., 2018; H. Wang, Ge, Uchida, Luu, & Ahn, 2011), is reduced and re-introducing Gli3R rescues the decreased formation of basal progenitors. Taken together, these findings implicate the primary cilium in controlling the decision of RGCs to either undergo direct neurogenesis or to form basal progenitors, thereby governing the neuronal subtype composition of the cerebral cortex.

## RESULTS

### *Inpp5e*^Δ/Δ^ embryos show mild telencephalic patterning defects

Controlling the balance between direct and indirect neurogenesis in the developing cerebral cortex is mediated by cell-cell signaling (Cardenas et al., 2018) and is hence likely to involve the primary cilium. To investigate potential ciliary roles, we started characterizing cortical stem cell development in embryos mutant for the *Inpp5e* gene which has a prominent role in ciliary signaling and stability. Mutations in ciliary genes have previously been found to result in telencephalon patterning defects, most notably in a ventralisation of the dorsal telencephalon and/or in defects at the corticoseptal (CSB) and pallial/subpallial boundaries (PSPB) (Ashique et al., 2009; Besse et al., 2011; Willaredt et al., 2008). Therefore, we first considered the possibility that such early patterning defects may be present in *Inpp5e* mutant embryos and could affect cortical stem cell development. In situ hybridization and immunofluorescence analyses of E12.5 control and *Inpp5e*^Δ/Δ^ embryos revealed no obvious effect on the expression of dorsal and ventral telencephalic markers at the corticoseptal boundary (SupFig. 1). In contrast, the pallial/subpallial boundary was not well defined with a few scattered Pax6+ and *Dlx2* expressing cells on the wrong side of the boundary, i.e. in the subpallium and pallium, respectively (SupFig. 1). Moreover, the hippocampal anlage appeared smaller and disorganized with low level and diffuse expression of cortical hem markers (SupFig. 2), consistent with known roles of Wnt/β-catenin and Bmp signaling in hippocampal development (Galceran, Miyashita-Lin, Devaney, Rubenstein, & Grosschedl, 2000; Lee, Tole, Grove, & McMahon, 2000). In contrast, progenitors in the neocortical ventricular zone of *Inpp5e* mutant mice expressed the progenitor markers *Emx1*, *Lhx2*, *Pax6* and *Ngn2,* though the levels of Pax6 protein expression appeared reduced in the medial neocortex suggestive of a steeper lateral to medial Pax6 expression gradient in mutant embryos. These expression patterns were maintained in E14.5 *Inpp5e*^Δ/Δ^ embryos but revealed an area in the very caudal/dorsal telencephalon where the neocortex was folded (SupFig. 3). At this level, folds were also present in the hippocampal anlage. Taken together, these findings indicate that *Inpp5e* mutants have mild patterning defects affecting the integrity of the PSPB, hippocampal development and the caudal-most neocortex while the rostral neocortex shows no obvious malformation and can therefore be analysed for effects of the *Inpp5e* mutation on direct and indirect neurogenesis.

### *Inpp5e* controls direct vs indirect neurogenesis in the lateral neocortex

Based upon these findings, we started analyzing the proliferation and differentiation of radial glial cells in *Inpp5e*^Δ/Δ^ embryos in the rostrolateral and rostromedial neocortex to avoid the regionalization defects described above. As a first step, we determined the proportion of radial glial cells, basal progenitors and neurons in these regions in E12.5 embryos. Double immunofluorescence for PCNA which labels all progenitor cells (Hall et al., 1990) and the radial glial marker Pax6 did not reveal differences in the proportions of radial glial cells at both medial and lateral levels (Fig. 1A-D). In contrast, the proportion of Tbr2+ basal progenitors was reduced laterally but not medially (Fig. 1E-H). This decrease coincided with an increase in Tbr1+ neurons specifically in the lateral neocortex (Fig. 1I-L). To determine whether these alterations are maintained at a later developmental stage, we repeated this investigation in E14.5 embryos. This analysis revealed no significant differences in the proportion of Pax6+ RGCs (Fig. 2A-D). Similarly, there was no alteration in the proportion of Tbr2+ basal progenitors in lateral neocortex, however, their proportion was reduced medially (Fig. 2E-H).

**Figure 1:**
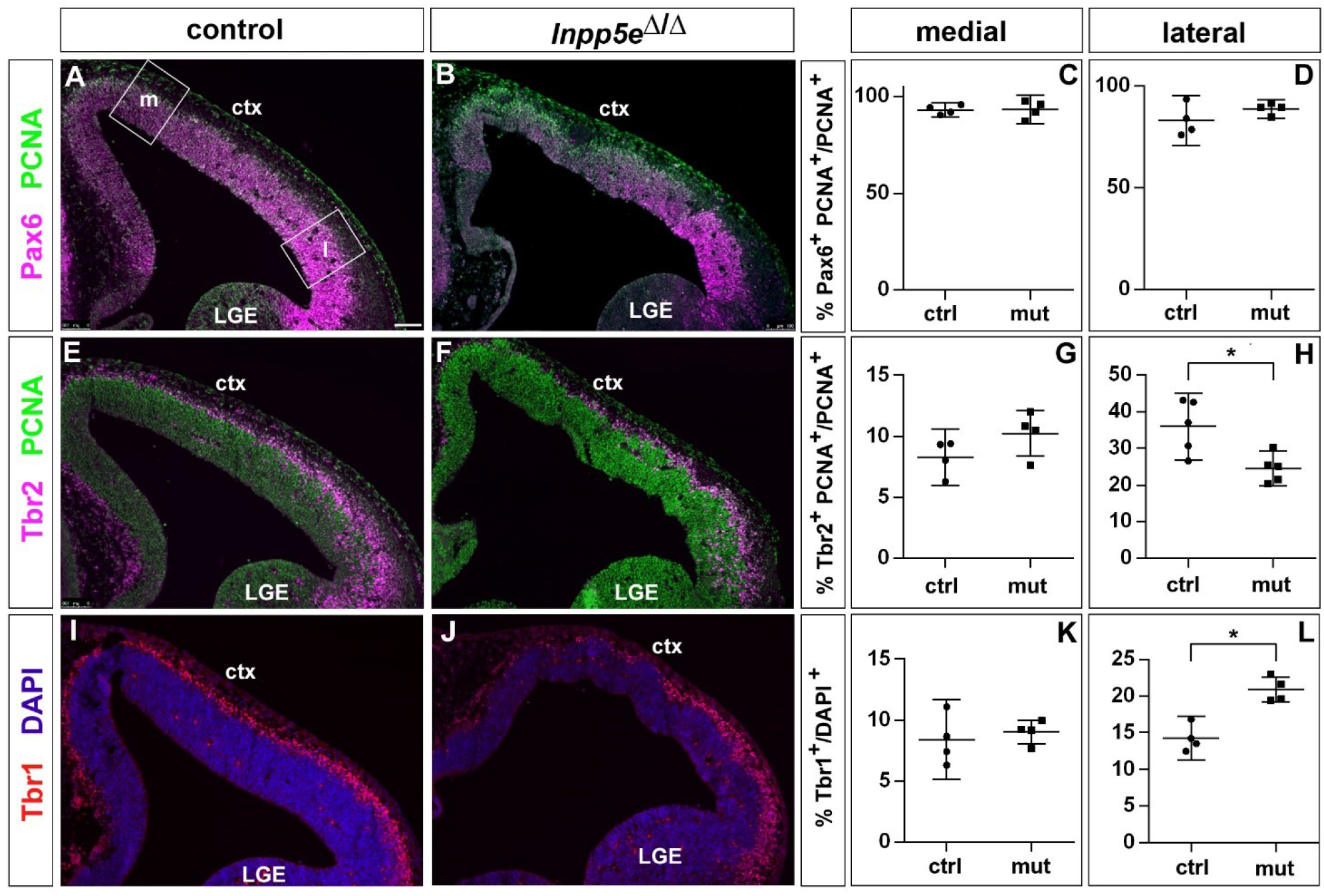
Increased neuron formation in the dorsolateral telencephalon of E12.5 *Inpp5*^Δ/Δ^ mutant embryos. (A-D) Pax6/PCNA double immunofluorescence staining revealed the proportion of apical radial glial cells which remained unaltered in the mutant. The boxes in (A) indicate the regions in the medial (m) and lateral (l) telencephalon at which cell counts were performed. (E-H) Reduced proportions of basal progenitors in the lateral telencephalon as revealed by staining for Tbr2 and PCNA. (I-L) Tbr1 immunostaining showed that the proportion of neurons is increased in the lateral telencephalon. All statistical data are presented as means ± 95% confidence intervals (CI); Mann Whitney tests; n = 4 except for (H) with n=5; * p < 0.05. Scale bar: 100μm. ctx: cortex; LGE: lateral ganglionic eminence.

**Figure 2:**
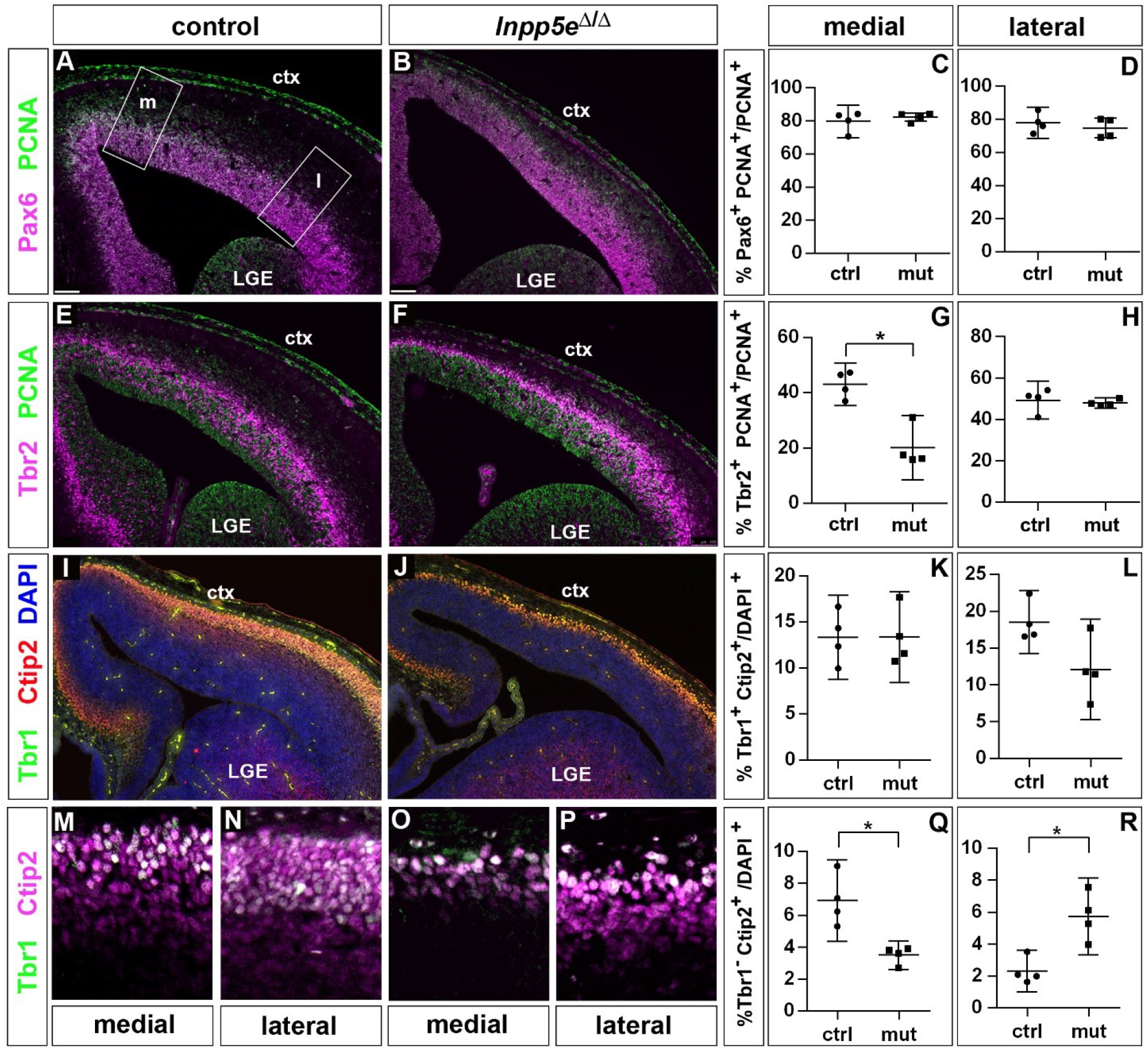
Proportions of radial glial cells, basal progenitors and neurons in the neocortex of E14.5 *Inpp5e*^Δ/Δ^ embryos. (A-D) The proportion of radial glial cells remains unaffected by the *Inpp5e* mutation as revealed by Pax6/PCNA double immunofluorescence. The boxes in A indicate the regions in the medial (m) and lateral (l) telencephalon at which cell counts were performed. (E-H) Tbr2/PCNA double staining showed a reduced proportion of basal progenitors in the *Inpp5e*^Δ/Δ^ medial but not lateral neocortex. (I-R) Neuron formation in the *Inpp5e*^Δ/Δ^ mutant necocortex. The proportion of Tbr1^+^ Ctip2^+^ neurons is not significantly altered (I-L) whereas the proportion of Tbr1^-^ Ctip2^+^ neurons is decreased and increased in the medial and lateral neocortex, respectively (M-R). All statistical data are presented as means ± 95% confidence intervals (CI); Mann Whitney tests; n = 4; * p < 0.05. Scale bars: 100μm (A) and 50μm (M). ctx: cortex; LGE: lateral ganglionic eminence.

To label cortical projection neurons, we used double immunofluorescence for Tbr1 and Ctip2 which allowed us to distinguish between Ctip2+Tbr1+ and Ctip2+Tbr1-neurons. Quantifying these subpopulations showed no effect on the formation of Ctip2+Tbr1+ neurons in *Inpp5e*^Δ/Δ^ embryos. In contrast, the proportion of Ctip2+Tbr1-neurons was reduced medially but increased in the lateral neocortex (Fig. 2I-R). Taken together, these findings show that initially Tbr1+ and later Ctip2+Tbr1-neurons were increasingly formed in the lateral neocortex of *Inpp5e*^Δ/Δ^ embryos and that the proportion of basal progenitors recovered after an initial down-regulation.

To address the defective cellular processes underlying these neurogenesis defects in *Inpp5e* mutants, we first measured proliferation rates of cortical progenitors and performed double immunofluorescence for PCNA and pHH3 which labels mitotic radial glial cells located at the ventricular surface and dividing basal progenitors in abventricular positions (SupFig. 4). This analysis revealed no statistically significant differences in the E12.5 and E14.5 lateral neocortex of control and *Inpp5e*^Δ/Δ^ embryos. The proportion of mitotic basal progenitors, however, was reduced in the E12.5 medial neocortex (SupFig. 4).

The cell cycle represents another key regulator of neuronal differentiation and a mutation in *Kif3a* affects ciliogenesis and the cell cycle in the developing neocortex (Wilson, Wilson, Wang, Wang, & McConnell, 2012). To investigate the possibility of altered cell cycle kinetics, we used a BrdU/IdU double labelling strategy (Martynoga, Morrison, Price, & Mason, 2005; Nowakowski, Lewin, & Miller, 1989) to determine S phase length and total cell cycle length in radial glial cells and found no statistically significant changes in these parameters (SupFig. 5).

Finally, the increased neuron production could also be explained by an increase in direct neurogenesis at the expense of basal progenitor cell formation. To test this possibility, we gave BrdU to E11.5 pregnant mice 24h before dissecting the embryos. We then used BrdU immunostaining in conjunction with Tbr1 and Tbr2 to identify the neurons and basal progenitors formed in the lateral neocortex within the 24h time period. This analysis showed that the proportion of Tbr1+ neurons compared to the total number of BrdU+ cells increased while the Tbr2+ proportion decreased in *Inpp5e* mutants (Fig. 3). Since the cell cycle of basal progenitors is longer than 24h (Arai et al., 2011), the 24h interval used in our cell cycle exit experiment was too short for newly formed basal progenitors to undergo one additional round of the cell cycle and as the BrdU label would have been diluted with a further round of division, this analysis supports our hypothesis that direct neurogenesis became more prevalent in *Inpp5e*^Δ/Δ^ radial glial cells.

**Figure 3:**
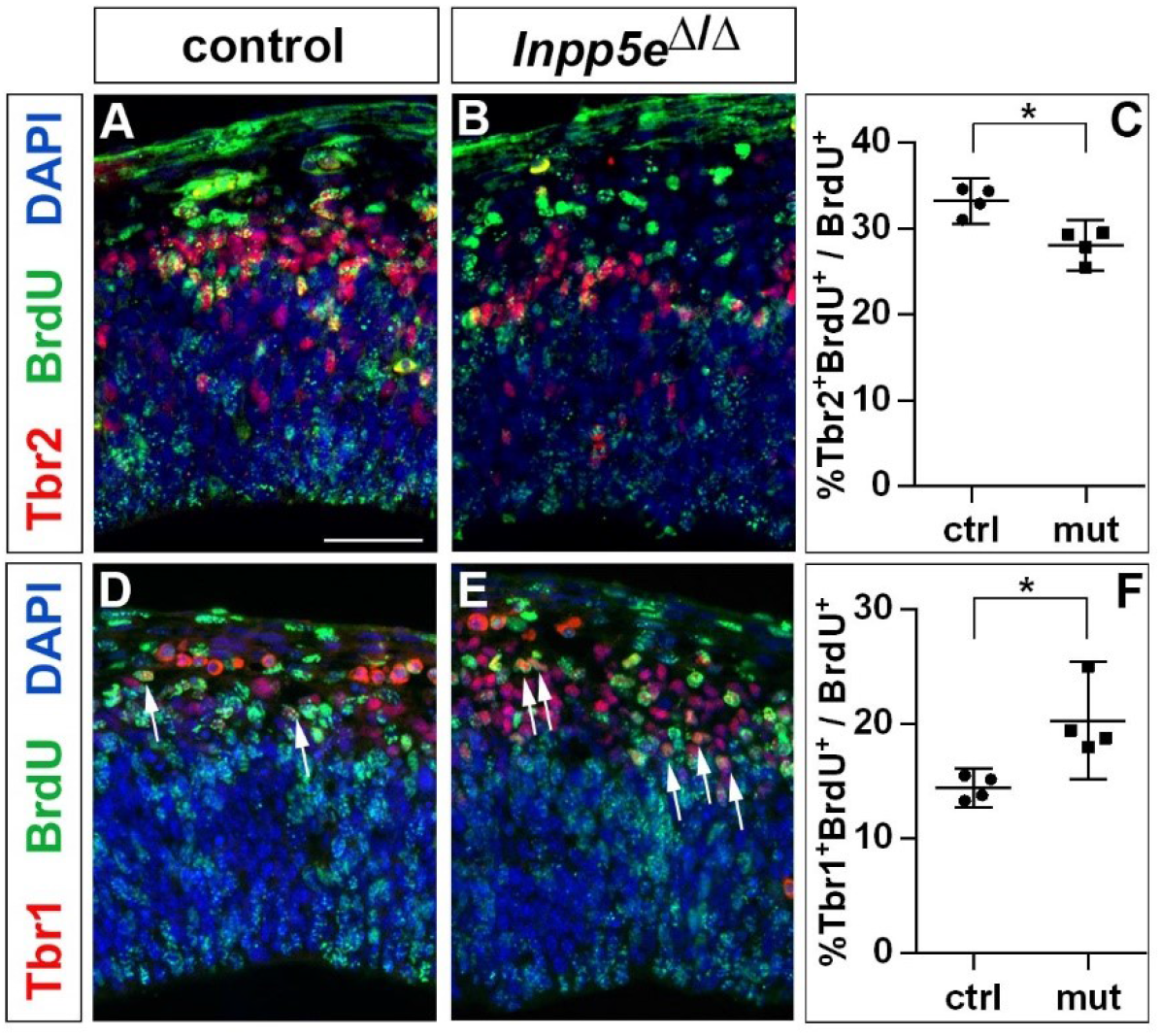
Increased neurogenesis at the expense of basal progenitor formation in E12.5 *Inpp5e* ^Δ/Δ^ mutants. Immunohistochemistry on sections of E12.5 control (A, D) and *Inpp5e*^Δ/Δ^ embryos (B, E) that were treated with BrdU 24 hours earlier. (A-C) Tbr2/BrdU double labelling showed that less basal progenitors formed from the BrdU labelled progenitor cohort. (D-F) The proportion of newly formed Tbr1^+^ neurons is increased. The arrows in D and E label Tbr1^+^ BrdU^+^ cells. All statistical data are presented as means ± 95% confidence intervals (CI); Mann Whitney tests; n = 4; * p < 0.05. Scale bar: 50μm.

### Cortical malformations in *Inpp5e*^Δ/Δ^ embryos

Next, we investigated the consequences of this increase in direct neurogenesis on cortical size and layer formation. Since *Inpp5e*^Δ/Δ^ newborn pups die perinatally (Bielas et al., 2009), we focused our analysis on E18.5 embryos. The mutant lacked obvious olfactory bulbs, as revealed by whole mounts of control and mutant brains (SupFig. 6). To gain insights into the overall histology of the mutant forebrain, we stained coronal sections with DAPI. This analysis showed that the mutant cortex was thinner laterally but not medially with a more pronounced reduction of the thickness at caudal levels (SupFig. 7). In addition, the hippocampus was malformed with a smaller dentate gyrus. Investigating the expression of markers characteristic of the entire hippocampus (*Nrp2*; (Galceran et al., 2000)), the CA1 field (Scip1; (Frantz, Bohner, Akers, & McConnell, 1994)) and the dentate gyrus (Prox1; (Oliver et al., 1993)) showed that these hippocampal structures were present but were severely reduced in size and disorganized in *Inpp5e*^Δ/Δ^ embryos (SupFig. 8). In addition, the corpus callosum, the major axon tract connecting the two cerebral hemispheres, was correctly formed but was smaller. We confirmed this effect by staining callosal axons and surrounding glial cells that guide these axons to the contralateral hemisphere with L1 and GFAP, respectively (SupFig. 9).

After characterizing the gross morphology of the *Inpp5e*^Δ/Δ^ cortex, we next investigated whether the increased neuron formation in E12.5 mutant embryos led to changes in the neuronal subtype composition of the E18.5 cortex. To this end, we used immunofluorescence labelling for Tbr1 and Ctip2 to analyse the formation of layer VI and V neurons, respectively, whereas Satb2 served as a layer II-IV marker (Fig. 4). Inspecting these immunostainings at low magnification showed that Tbr1+, Ctip2+ and Satb2+ neurons occupied their correct relative laminar positions in *Inpp5e* mutants (Fig. 4A-F) except for neuronal heterotopias which were present in all mutant brains, though their number and position varied (Fig. 4D). These immunostainings also revealed a medial shift in the position of the rhinal fissure, a sulcus that is conserved across mammalian species and separates neocortex from the paleocortical piriform cortex (Ariens-Kapers, Huber, & Crosby, 1936). This shift was more marked caudally and suggests a dramatic expansion of the *Inpp5e* mutant piriform cortex at the expense of neocortex at caudal most levels (Fig. 4D-F). Using the Tbr1/Ctip2 and Satb2 stainings, we determined the proportions of deep and superficial layer neurons, respectively. Because of the expanded piriform cortex in *Inpp5e* mutants, we limited this investigation to the unaffected rostral neocortex. In the rostrolateral neocortex, we found the proportion of Tbr1+ neurons to be reduced (Fig. 4G, H, M). This reduction coincided with an increased proportion of Ctip2+ layer V neurons (Fig. 4I, J, N) while the Satb2 population was unchanged (Fig. 4K, L, O). In contrast, the rostromedial neocortex did not show any differences (Fig. 4P-X). Thus, the increase in direct neurogenesis in the lateral neocortex during earlier development concurs with a change in the proportions of E18.5 Tbr1+ and Ctip2+ deep layer neurons.

**Figure 4:**
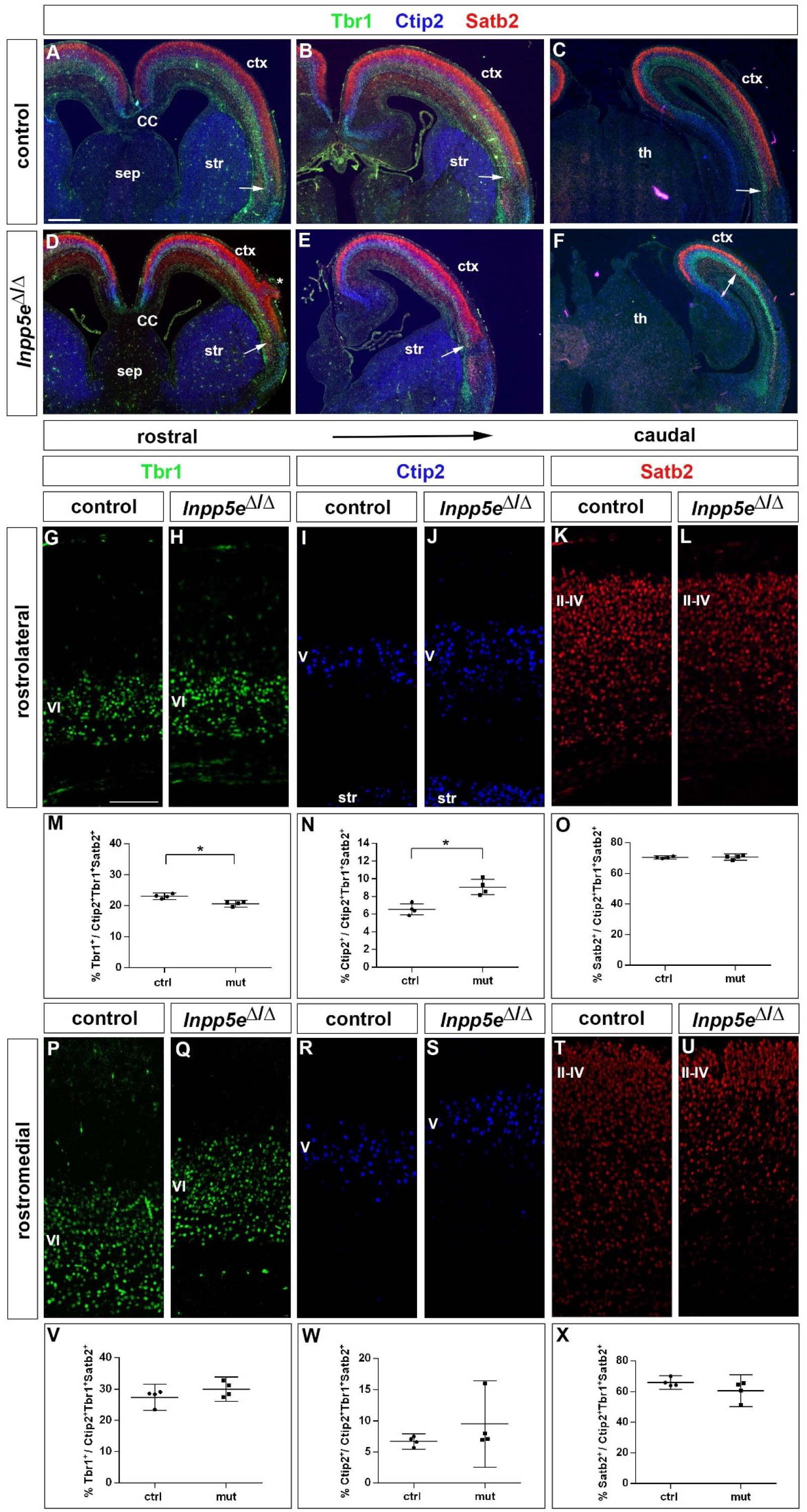
Increased formation of layer V neurons in E18.5 *Inpp5e* ^Δ/Δ^ mutants. (A-F) Coronal sections immunostained for the deep layer markers Tbr1 (layer VI) and Ctip2 (layer V) and for the upper layer marker Satb2 (layers II-IV); there is no obvious defect in layering in the *Inpp5e*^Δ/Δ^ mutant except for the formation of a heterotopia (asterisk in D). At caudal levels, the cortex becomes thinner and the rhinal fissure is shifted medially as indicated by the arrows. (G-O) Formation of cortical neurons at rostrolateral levels. The proportion of Tbr1+layer VI neurons is decreased with a concomitant increase in Ctip2+layer V neurons. (P-X) Portion of cortical neurons at rostromedial levels. Immunolabeling with cortical layer markers revealed no significant difference. Note that due to the thinner cortex, the position of layer VI Tbr1+ (Q) and layer V Ctip2+ neurons (J, S) appears to be shifted to more superficial positions, however, the relative order of these layers remains unaffected. All statistical data are presented as means ± 95% confidence intervals (CI); Mann Whitney tests; n = 4; * p < 0.05. Scale bars: 500μm (A) and 100μm (G). CC: corpus callosum; ctx: cortex; sep: septum; str: striatum.

### A mutation in the ciliary gene *Tctn2* leads to increased telencephalic neurogenesis

To start to unravel the mechanisms by which *Inpp5e* controls cortical stem cell development, we first analysed whether the increased early neurogenesis is restricted to *Inpp5e*^Δ/Δ^ mutants or is observed in another mutant affecting cilia. To this end, we focused on the *TECTONIC* 2 (*TCTN2*) gene which is crucial for ciliary transition zone architecture (Shi et al., 2017) and which, like *INPP5E*, is mutated in Joubert Syndrome (Garcia-Gonzalo et al., 2011). Interestingly, E12.5 *Tctn2*^*−/−*^ mutant embryos (Reiter & Skarnes, 2006) also showed an increased proportion of Tbr1+ projection neurons and a concomitant decrease in Tbr2+ basal progenitors in the dorsolateral telencephalon (Fig. 5). Due to embryonic lethality, however, we were not able to investigate the formation of cortical neurons at later stages.

**Figure 5:**
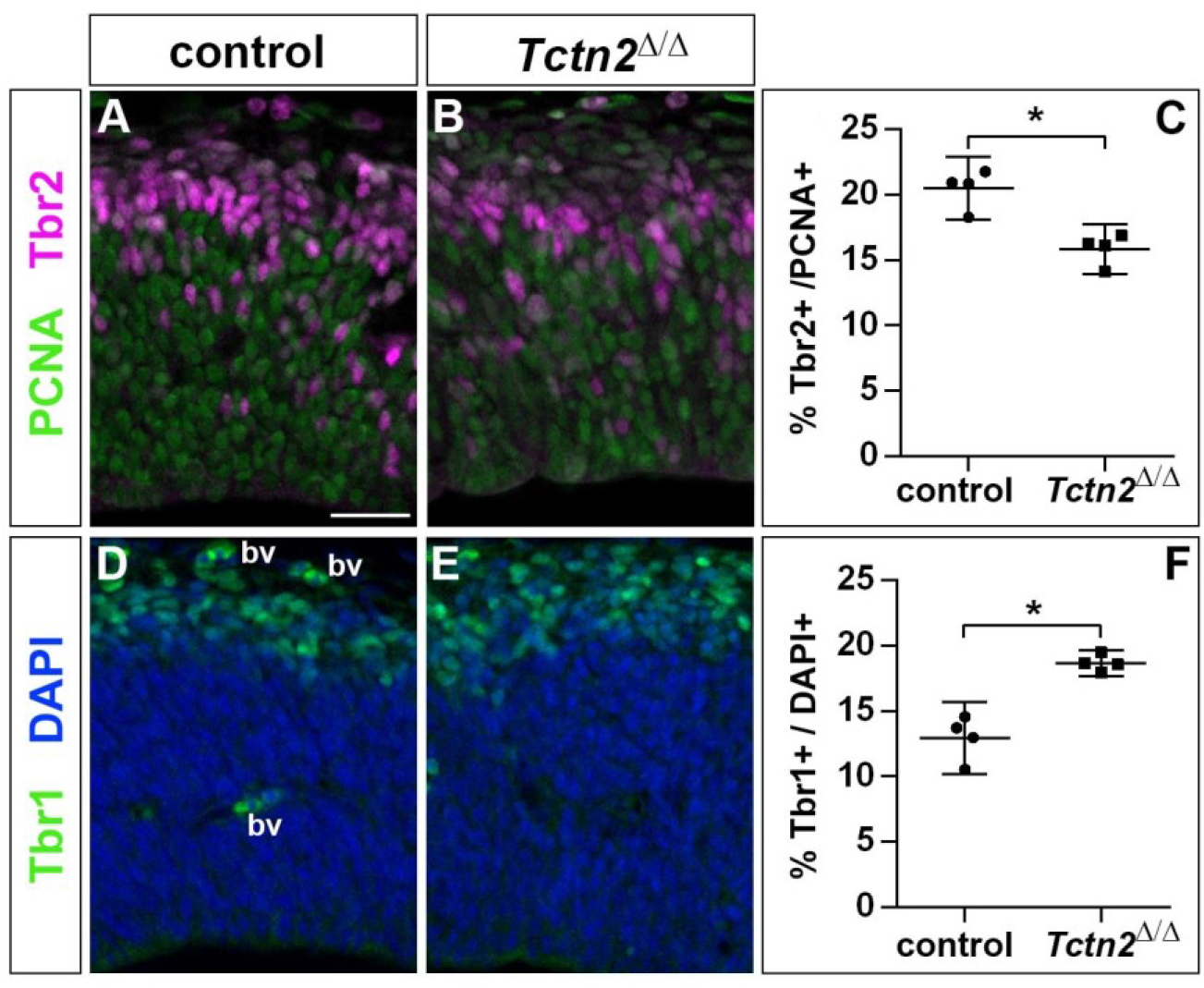
Increased generation of cortical neurons in E12.5 *Tctn2*^−/−^ embryos. (A-C) Double immunofluorescence for PCNA and Tbr2 revealed a significantly decreased proportion of basal progenitors. (D-F) The portion of Tbr1^+^ cortical neurons is increased. All statistical data are presented as means ± 95% confidence intervals (CI); Mann Whitney tests; n = 4; * p < 0.05. Scale bar: 50μm. bv: blood vessel.

### Ciliary defects in the forebrain of E12.5 *Inpp5e*^Δ/Δ^ embryos

Our findings in the *Inpp5e* and *Tctn2* mutants suggested a role for cilia in cortical progenitor cells to control early neurogenesis. Therefore, we examined the presence and the structure of primary cilia in the developing forebrain of *Inpp5e*^Δ/Δ^ embryos by immunofluorescence and electron microscopy. We first analyzed the presence of the small GTPase Arl13b, enriched in ciliary membranes, and of *γ*-Tubulin (*γ*Tub), a component of basal bodies (Caspary, Larkins, & Anderson, 2007). We found no major difference in the number, the apical localization or the size of cilia in control and *Inpp5e*^Δ/Δ^ neuroepithelial cells in the E12.5 telencephalon (Fig. 6A,B) or diencephalon (data not shown).

**Figure 6:**
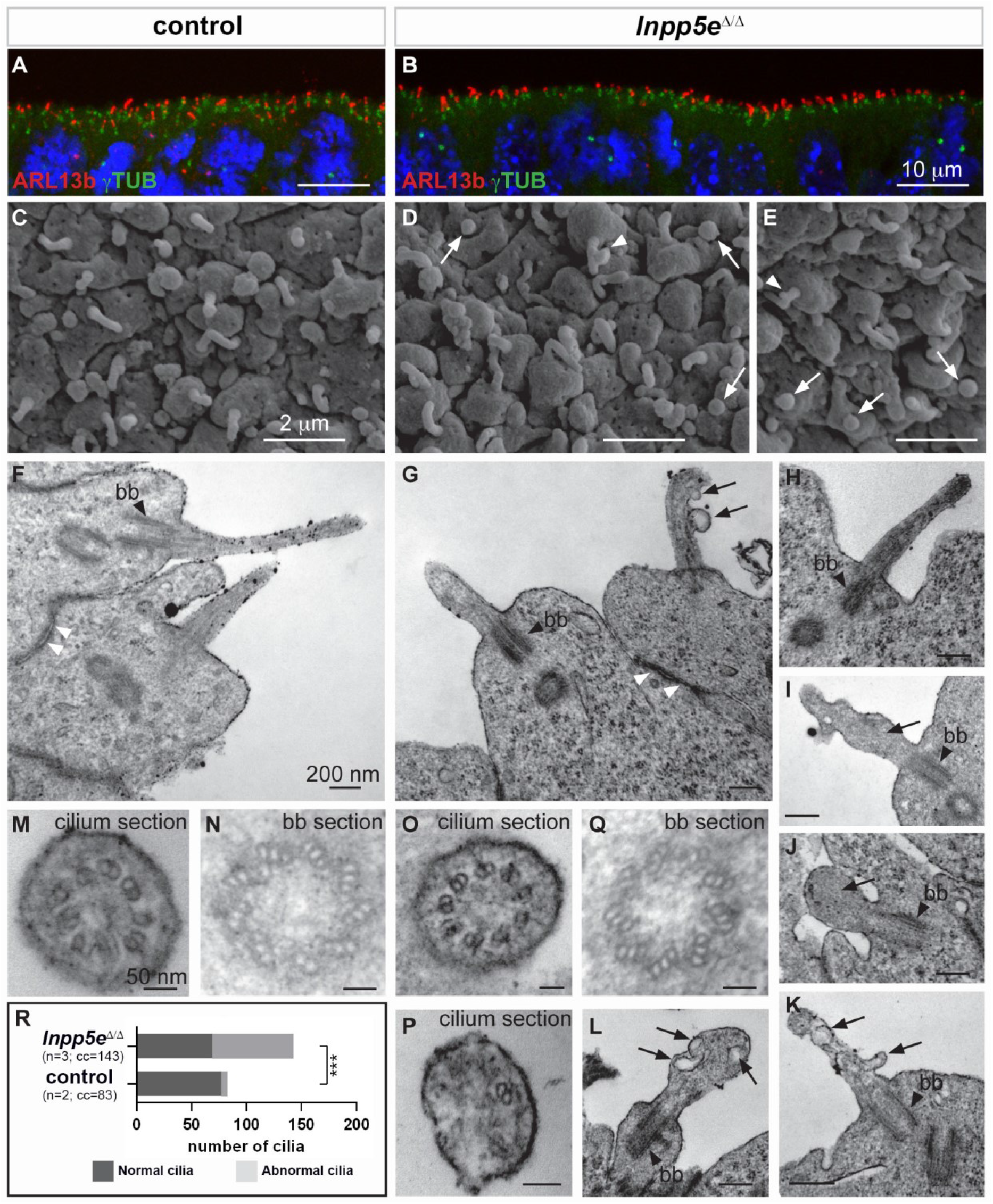
Ciliary defects in E12.5 *Inpp5e* ^Δ/Δ^ forebrain. (A-B) Immunohistochemistry for Arl13b and *γ*-Tubulin (*γ*TUB) on E12.5 brain cryosections showed an accumulation of ciliary axonemes and basal bodies, respectively, at the apical border of radial glial cells facing the ventricules in both control (A) and *Inpp5e*^Δ/Δ^ (B) embryos without any obvious difference. Scale bars: 10 μm. (C-E) Scanning electron microscopy (SEM) on E12.5 control (C) and *Inpp5e*^Δ/Δ^ (D, E) brains highlighted the presence of primary cilia projecting from the apical surface of radial glial cells in both control (A) and *Inpp5e*^Δ/Δ^ (B) embryos. However, SEM also revealed the presence of abnormal cilia in *Inpp5e*^Δ/Δ^ embryos having a spherical shape (arrows in D and E) or aberrant lateral buddings (arrowheads in D and E). Scale bars: 2 μm. (F-L) Transmission electron microscopy (TEM) analysis on E12.5 brains showed longitudinal sections of primary cilia in control (F) and *Inpp5e*^Δ/Δ^ (G-L) embryos. In control primary cilia, the axoneme appeared as an extension of the basal body (bb, black arrowheards) (F-H). In addition to cilia with normal morphology, abnormal cilia were identified in *Inpp5e*^Δ/Δ^ embryos thanks to the presence of a basal body apparently correctly docked to the apical membrane. Abnormal cilia lacked an axoneme (I, J, L) or showed unusual membranous structures, such as budding (G, K) or internal (I, K, L) vesicles (arrows) or undulating peripheral membranes (I). Note that tight junctions (white arrowheads in F and G) appeared normal in *Inpp5e*^Δ/Δ^ (G) and control (F) embryos, suggesting that apico-basal polarity of *Inpp5e*^Δ/Δ^ radial glial cells was not compromised. Scale bars: 200 nm. (M-Q) TEM images showing transverse sections of the axoneme (M, O, P) and the basal body (N, Q) in control (M, N) and *Inpp5e*^Δ/Δ^ (O-Q) embryos with no major difference in the basal bodies between control (N) and *Inpp5e*^Δ/Δ^ (Q) embryos. However, transverse section of primary cilia in *Inpp5e*^Δ/Δ^ brains revealed the presence of normal axonemes composed of 9 correctly organized doublets of microtubules on some radial glial cells (O), while others harboured an abnormal axoneme containing a lower number of microtubule doublets (P). Scale bars: 50 nm. (R) Graph showing the number of normal versus abnormal cilia (cil.) found on TEM images from control (n=3) or *Inpp5e*^Δ/Δ^ (n=3) embryos. cc: counted cilia.

To gain insights into the fine structure of these primary cilia we performed electron microscopy analyses. Scanning electron microscopy (SEM) provided an observation of the cilia protruding into the telencephalic ventricles. In control embryos, almost all radial glial cells had a single, ~1 μm long primary cilium (Fig. 6C), as previously described (Besse et al., 2011). Some *Inpp5e*^Δ/Δ^ mutant cells also displayed an apparently normal cilium (Fig. 6D, E), whereas other cells harbored abnormal cilia, either with a lateral blob (arrowhead in Fig. 6D) or as a short and bloated cilium-like protrusion (arrows in Fig. 6D,E).

Transmission electron microscopy (TEM) confirmed the presence of abnormal cilia in *Inpp5e*^Δ/Δ^ embryos. Cilia were recognized by basal bodies anchored to the apical membrane in both control and *Inpp5e*^Δ/Δ^ radial glial cells (Fig. 6F-L, N, Q). However, in *Inpp5e*^Δ/Δ^ cells, some cilia lacked the axoneme and showed unusual membranous structures that resemble budding vesicles emerging from the lateral surface of the cilium (Fig. 6G, K), internal vesicles (arrows in Fig. 6I, K, L), or undulating peripheral membranes (Fig. 6I), indicating an *Inpp5e*-dependent defect in ciliary membrane morphology. Transverse sections revealed the presence of cilia with apparently normal 9+0 axonemes, as well as cilia containing abnormal numbers of microtubule doublets in *Inpp5e*^Δ/Δ^ embryos (Fig. 6O, P). To quantify these ciliary defects, we counted the number of normal versus abnormal cilia on TEM images obtained from control and *Inpp5e*^Δ/Δ^ embryos, and found an increase in abnormal cilia in *Inpp5e*^Δ/Δ^ compared to control embryos (Fig. 6R). Taken together, a significant number of abnormal primary cilia were found at the apical end of E12.5 radial glial cells in the forebrain of *Inpp5e*^Δ/Δ^ embryos. These abnormalities are consistent with a role of *Inpp5e* in maintaining cilia stability (Jacoby et al., 2009).

### Receptor tyrosine kinase and mTOR signaling in *Inpp5e*^Δ/Δ^ embryos

Given these defects in ciliary structure, we started to investigate cilia-controlled signaling pathways, defects in which might underlie the increased direct neurogenesis in the *Inpp5e*^Δ/Δ^ mutant radial glial cells. Inpp5e hydrolyzes the 5-phosphate of PI(3,4,5)P_3_ which serves as a second messenger in receptor tyrosine kinase signaling. We therefore investigated the amount and distribution of pErk1/2 as a read-out of receptor tyrosine kinase signaling. Western blots using dissected dorsal telencephalon from E12.5 and E14.5 embryos revealed no difference in the ratio between pErk1/2 and Erk1/2 (SupFig. 10). Moreover, pErk1/2 expression in the E12.5 forebrain remains restricted to the LGE, the diencephalon and to meninges covering the telencephalon but was not identified in cortical progenitors (SupFig. 10). In E14.5 embryos, we detected pErk1/2 immunoreactivity in cortical progenitors in a lateral to medial gradient covering the whole extent of the neocortex. In contrast, high level pErk1/2 expression remained confined to the lateral neocortex with little or no expression medially in *Inpp5e*^Δ/Δ^ embryos (SupFig. 10). These findings suggest that pErk signaling does not play a role in the early neurogenesis defect in *Inpp5e* mutants but may be involved in the recovery of basal progenitor formation.

PI(3,4,5)P_3_, a substrate of the Inpp5e phosphatase, is essential for the effective activation of the serine threonine kinase Akt (Kisseleva, Cao, & Majerus, 2002; Plotnikova et al., 2015) raising the possibility that loss of *Inpp5e* results in activated Akt signaling in the neocortex. Moreover, phosphorylation of Akt at Serine^473^ has been implicated in inhibiting cilia assembly and promoting cilia disassembly (Mao et al., 2019). We therefore investigated the amount of phospho-Akt^S473^ in western blots on protein extracts from the E12.5 and E14.5 dorsal telencephalon. This analysis revealed increased phosphorylation of Akt at Serine^473^ at E12.5 but no significant change at E14.5 (SupFig. 11). Since Akt acts upstream of mTOR (Yu & Cui, 2016), we also analysed the activity of the mTOR pathway and investigated the phosphorylation of the S6 ribosomal protein, at Serine^235/236^ and Serine^S240/244^ (Ferrari, Bandi, Hofsteenge, Bussian, & Thomas, 1991). Interestingly, *Inpp5e* mutants showed increased S6 phosphorylation at Serine^235/236^ in E12.5 and E14.5 embryos whereas an increase in phospho-S6^S240/244^ remained restricted to E12.5 (SupFig. 11). Taken together, these findings indicate that *Inpp5e* inactivation leads to increased activation of Akt and mTOR signaling in the E12.5 dorsal telencephalon.

Next, we investigated a potential role of elevated mTOR signaling in controlling the balance between direct and indirect neurogenesis and treated E11.5 pregnant mice with the mTORC1 inhibitor rapamycin (1mg/kg bodyweight). We harvested embryos 24h later and determined the proportions of basal progenitors and neurons. This analysis revealed that administering rapamycin did not rescue basal progenitor and neuron formation although the cortex of rapamycin treated *Inpp5e*^Δ/Δ^ embryos appeared elongated (SupFig. 11). Based on this experiment, augmented mTOR signaling does not underlie the altered ratio of direct vs indirect neurogenesis in *Inpp5e*^Δ/Δ^ embryos.

### Restoring Gli3 repressor ratio rescues cortical malformations in *Inpp5e*^Δ/Δ^ embryos

Primary cilia also play a crucial role in Shh signaling by controlling the proteolytic cleavage of full length Gli3 (Gli3FL) into the Gli3 repressor form (Gli3R) in the absence of Shh and by converting Gli3FL into the transcriptional activator Gli3A in the presence of Shh. Moreover, the dorsal telencephalon predominately forms Gli3R (Fotaki, Yu, Zaki, Mason, & Price, 2006) and mice that can only produce Gli3R have no obvious defect in cortical development (Besse et al., 2011; Böse, Grotewold, & Rüther, 2002). In addition, we recently showed that Gli3 has a prominent role in radial glial cells controlling the switch from symmetric proliferative to asymmetric neurogenic cell division (Hasenpusch-Theil et al., 2018). Therefore, we hypothesized that alterations in Gli3 processing caused by abnormal cilia function underlies the increased direct neurogenesis and the cortical malformations in *Inpp5e*^Δ/Δ^ embryos. In situ hybridization showed that *Gli3* mRNA expression might be slightly reduced but the overall expression pattern in the telencephalon remains unaffected (SupFig. 12). We next investigated the formation of Gli3FL and Gli3R in the E12.5 dorsal telencephalon of *Inpp5e*^Δ/Δ^ embryos using Western blots. This analysis revealed no change in the levels of Gli3FL but a significant decrease inGli3R which resulted in a reduced Gli3R to Gli3FL ratio in the mutant (Fig. 7A-D) suggesting that the *Inpp5e* mutation affects Gli3 processing.

**Figure 7:**
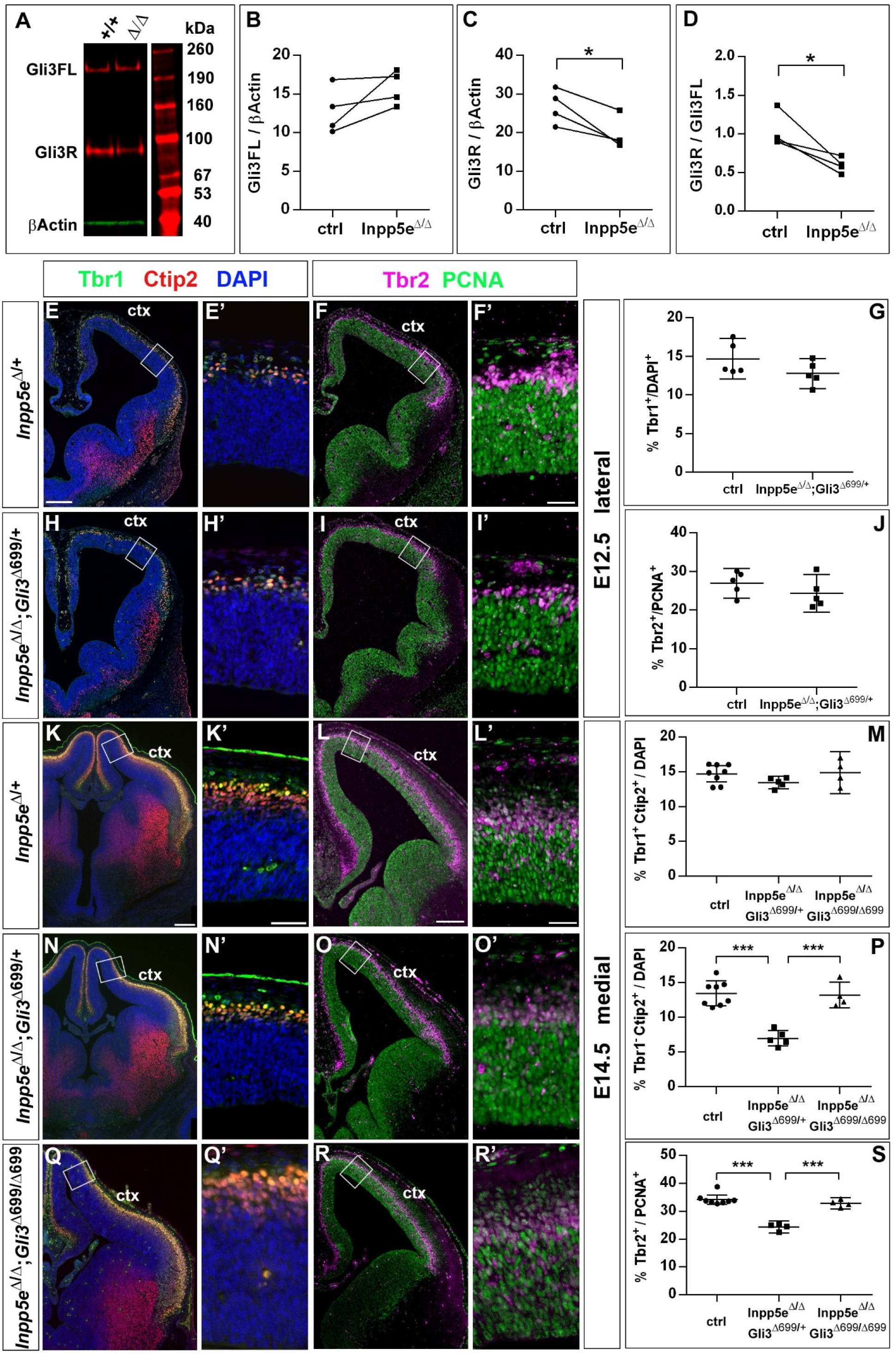
Re-introducing the Gli3 repressor rescues the neurogenesis defect in *Inpp5e* mutants. (A-D). Gli3 Western blot on E12.5 dorsal telencephalic tissue revealed the Gli3 full length (FL) and repressor (R) forms (A). While Gli3FL levels are not affected (B), levels of Gli3R (C) and the Gli3R/Gli3FL ratio (D) are decreased in *Inpp5e*^Δ/Δ^ embryos. A paired t-test was used to evaluate levels of Gli3FL and Gli3R and the Gli3R/Gli3FL ratio in four control/*Inpp5e*^Δ/Δ^ embryo pairs derived from four different litters. (E-S) Formation of basal progenitors and neurons in the neocortex of *Inpp5e*^Δ/Δ^;Gli3^Δ699/+^ and *Inpp5e*^Δ/Δ^Gli3^Δ699/699^ embryos. (E-N) In the lateral neocortex of E12.5 embryos, there is no significant difference in the proportions of Tbr1+ neurons (E-G) and basal progenitor cells (H-J) between control and *Inpp5e*^Δ/Δ^;Gli3^Δ699/+^ embryos. (K-S) In the E14.5 medial neocortex, the proportion of Tbr1^-^ Ctip2^+^ neurons (P) and basal progenitors (S) are significantly reduced in *Inpp5e*^Δ/Δ^;*Gli3*^Δ699/+^ embryos but not in *Inpp5e*^Δ/Δ^;Gli3^Δ699/699^ embryos. All statistical data except for the Western blot data (C, D) are presented as means ± 95% confidence intervals (CI); Mann Whitney (n=5) (G, J) and ONE way ANOVA followed by Tukey’s multiple comparison test (J, M, P); * p < 0.05; ** p < 0.01; *** p < 0.001. Scale bars: 50μm (E, K, L), 100μm (A), and 250μm (F’, K’, L’). ctx: cortex; MGE: medial ganglionic eminence; LGE: lateral ganglionic eminence.

The next set of experiments aimed to clarify a role for the reduced Gli3 processing. To this end, we restored Gli3R levels by crossing *Inpp5e* mutants with *Gli3*^Δ699/+^ mice that can only produce Gli3R in a cilia-independent manner (Besse et al., 2011; Böse et al., 2002). Overall inspection of *Inpp5e*^Δ/Δ^*Gli3*^Δ699/+^ embryos revealed normal eye formation whereas *Inpp5e*^Δ/Δ^ embryos either completely lacked eyes or showed microphthalmia (Jacoby et al., 2009) (SupFig. 13). Moreover, the overall morphology of the telencephalon is much improved in *Inpp5e*^Δ/Δ^*Gli3*^Δ699/+^ embryos as compared to *Inpp5e*^Δ/Δ^ embryos. In E18.5 *Inpp5e*^Δ/Δ^*Gli3*^Δ699/+^ mutants, the corpus callosum has a thickness indistinguishable from that of control embryos (SupFig. 14). In E12.5 and E14.5 *Inpp5e*^Δ/Δ^*Gli3*^Δ699/+^ embryos, the neocortex lacks the undulations of the VZ present in *Inpp5e*^Δ/Δ^ embryos (data not shown) and the morphology of the hippocampal anlage is more akin to that in wild-type embryos but it is still smaller and less bulged (Fig. 7E, H, K, N).

We also determined the proportions of basal progenitors and Tbr1+ neurons at E12.5 which were decreased and increased, respectively, in the lateral neocortex of *Inpp5e*^Δ/Δ^ embryos. Remarkably, there was no statistically significant difference between control and *Inpp5e*^Δ/Δ^*Gli3*^Δ699/+^ embryos (Fig. 7E-J) indicating that the neurogenesis phenotype of E12.5 *Inpp5e*^Δ/Δ^ mutants is rescued by a single copy of the Gli3^Δ699^ allele. The proportion of Tbr1+Ctip2+ neurons was not affected in the medial neocortex of E14.5 *Inpp5e*^Δ/Δ^*Gli3*^Δ699/+^ embryos (Fig. 7K, M, N) either, however, the proportion of Tbr1-Ctip2+ neurons was reduced (Fig. 7K, N, P). Similarly, the proportions of basal progenitors in the E14.5 medial neocortex was reduced in *Inpp5e*^Δ/Δ^*Gli3*^Δ699/+^ embryos as in *Inpp5e*^Δ/Δ^ embryos (Fig. 7L, O, S). As re-introducing a single Gli3^Δ699^ allele does not completely rescue the *Inpp5e*^Δ/Δ^ neurogenesis phenotype, we generated *Inpp5e*^Δ/Δ^ embryos homozygous for Gli3^Δ699^. Interestingly, the morphology of the telencephalon including the hippocampal formation was indistinguishable between control and *Inpp5e*^Δ/Δ^*Gli3*^Δ699/Δ699^ embryos (Fig. 7K, Q) and the proportions of Tbr1-Ctip2+ and basal progenitors were not affected any longer (Fig. 7K, L. P-S). Taken together, these findings indicate that re-introducing a single copy of the *Gli3R* allele into the *Inpp5e* mutant background leads to a partial rescue of cortical neurogenesis in *Inpp5e*^Δ/Δ^ embryos whereas two copies are required for a full rescue.

## DISCUSSION

Generating a functional cerebral cortex requires a finely tuned balance between direct and indirect neurogenesis to form subtypes of cortical projection neurons in appropriate numbers. Here, we show that the ciliary mouse mutants *Inpp5e* and *Tctn2* present with a transient increase in neurons forming directly from radial glia progenitors in the lateral neocortex at the expense of basal progenitor formation. This increase in neurogenesis results in augmented formation of Ctip2+ layer V neurons in the *Inpp5e* mutant cortex. Our studies also revealed that the *Inpp5e* mutation interfered with the stability of the RGC primary cilium and its signaling functions, leading to a reduction in the Gli3R levels. Since re-introducing Gli3R in an *Inpp5e* mutant background restored the decreased formation of normal proportions of basal progenitors and neurons, our findings implicate a novel role for primary cilia in controlling the signaling events that direct the decision of RGCs to undergo either direct or indirect neurogenesis.

### Primary cilia affect the decision between direct and indirect neurogenesis

Radial glial cells in the developing mouse neocortex have the potential to undergo symmetric proliferative or asymmetric cell divisions with the latter division mode producing neurons in a direct manner or indirectly via basal progenitors. Balancing out these division modes is important not only to determine final neuronal output and cortical size but also the types of cortical projection neurons and, hence, subtype composition of the adult neocortex. In the E12.5 *Inpp5e* and *Tctn2* mouse mutants, we identified an increased formation of neurons in the lateral neocortex. Based on our cell cycle exit experiment additional neurons are formed from RGCs at the expense of basal progenitors. Given the cell cycle length of basal progenitors of >24 hours (Arai et al., 2011), it is unlikely that new born basal progenitors would have undergone an additional round of cell division to produce two neurons within the time frame of this experiment. Such an extra division would also have diluted the BrdU label. We therefore conclude that the *Inpp5e* mutation caused RGCs to preferentially produce neurons directly.

Interestingly, this increase in direct neurogenesis led to an increased proportion of Ctip2+ deep layer V neurons in the E18.5 neocortex but did not coincide with a reduced proportion of upper layer neurons. This effect could be explained in several mutually non-exclusive ways. First, neurons born at E12.5 initially express both Ctip2 and Tbr1 (Fig. 7) and later down-regulate Ctip2. *Inpp5e* could therefore affect the signaling that controls this downregulation. Secondly, the proportions of basal progenitors and neurons was normalized in E14.5 mutants. Since basal progenitors are a main source of upper layer neurons (Arnold et al., 2008; Vasistha et al., 2015), this normalization would account for the sufficient numbers of Satb2+ upper layer neurons. Newly formed projection neurons signal back to RGCS via Jag1, Fgf9 and Neurotrophin 3 (Parthasarathy et al., 2014; Seuntjens et al., 2009; W. Wang et al., 2016) to control the sequential production of deep and upper layer neurons and of glia (Silva et al., 2019). *Inpp5e* might affect these signals by controlling cilia stability and/or levels of PI(3,4,5)P_3_ (Bielas et al., 2009; Jacoby et al., 2009) that acts as a second messenger in receptor tyrosine kinase signaling. This possibility is supported by our observation that the extent of the pErk expression domain in the neocortex is diminished in E14.5 *Inpp5e*^Δ/Δ^ embryos. Regardless of the exact mechanism, our findings suggest a novel, spatially and temporally restricted role for *Inpp5e* in controlling the decision between direct and indirect neurogenesis. This function differs from those described for other cilia mutants. Conditional inactivation of *Ift88* and *Kif3a* leads to a larger cortex (Foerster et al., 2017; Wilson et al., 2012) with a modest increase in BP production in the absence of a delay in neurogenesis (Foerster et al., 2017) while *Rpgrip1l* mutants have reduced numbers of both basal progenitors and neurons (Postel, Karam, Pezeron, Schneider-Maunoury, & Clement, 2019). These findings highlight the multiple and varied roles cilia play in cortical development.

### *Inpp5e* controls direct/indirect neurogenesis through Gli3 processing

Our study also shed lights into the mechanisms by which *Inpp5e* controls the decision between direct/indirect neurogenesis. Most notably, the Gli3R level and Gli3R/Gli3FL ratio are decreased in *Inpp5e*^Δ/Δ^ embryos. While the *Inpp5e* mutation does not lead to an up-regulation of Shh signaling in the dorsal telencephalon (Magnani et al., 2015), re-introducing a single or two copies of Gli3R in an *Inpp5e* mutant background partially and fully restores the neurogenesis defects, respectively. This rescue indicates that reduced levels of Gli3R rather than the reduction in the Gli3R/Gli3FL ratio are responsible for the prevalence of direct neurogenesis in *Inpp5e*^Δ/Δ^ embryos. This idea is consistent with the findings that (i) *Gli3*^Δ699/Δ699^ embryos that cannot produce Gli3FL and Gli3A show no obvious phenotype in cortical development (Besse et al., 2011; Böse et al., 2002), (ii) dorsal telencephalic patterning defects in *Gli3*^Xt/Xt^ mutants are not rescued in *Shh*^−/−^/*Gli3*^XtXt^ double mutants (Rallu et al., 2002; Rash & Grove, 2007), (iii) Shh promotes the generation of olfactory bulb interneurons and cortical oligodendrocytes and neurogenesis in the subventricular zone by reducing Gli3R rather than by promoting Gli activator function (Petrova, Garcia, & Joyner, 2013; H. Wang, Kane, Lee, & Ahn, 2014; Zhang et al., 2020). In addition, there is also a dramatic rescue of eye development and the rescue also extends to other malformations of the *Inpp5e*^Δ/Δ^ forebrain, including the corpus callosum, the hippocampus and the expansion of the piriform cortex, structures that are also affected in *Gli3* null and hypomorphic mutants (Amaniti et al., 2015; Johnson, 1967; Magnani et al., 2014; Theil, Alvarez-Bolado, Walter, & Rüther, 1999; Wiegering, Petzsch, Kohrer, Ruther, & Gerhardt, 2019). Taken together, these findings support the idea that *Inpp5e* and the primary cilium control key processes in cortical development by regulating the formation of Gli3R.

Our analyses support several mutually non-exclusive mechanisms how the *Inpp5e* mutation impacts on Gli3 processing. First, our electron microscopy study revealed severe structural abnormalities in large proportions of cilia. The Inpp5e phosphatase hydrolyses PI(3,4,5)P_3_, which is essential for the effective activation of the serine threonine kinase Akt (Kisseleva et al., 2002; Plotnikova et al., 2015). Following PIP3 binding, Akt translocates to the membrane and becomes phosphorylated at T308 by phosphoinositide dependent kinase-1 (Pdk1) and at S473 by mammalian target of rapamycin complex (mTORC2) (Yu & Cui, 2016). Consistent with the loss of *Inpp5e* function and a resulting increase in PI(3,4,5)P_3_, our western blot analysis revealed elevated pAkt^S473^ levels. Increased phosphorylation at this site has been implicated in inhibiting cilia assembly and promoting cilia disassembly (Mao et al., 2019) and could hence explain the structural defects of RGC *Inpp5e*^Δ/Δ^ cilia. Secondly, *Inpp5e* could control Gli3 processing through its effect on the transition zone (TZ). It is required for TZ molecular organisation (Dyson et al., 2017) and its substrate PI(4,5)P2 plays a role in TZ maturation in Drosophila (Gupta, Fabian, & Brill, 2018). This model is further supported by our finding that a mouse mutant for the TZ protein Tctn2 phenocopies the *Inpp5e*^Δ/Δ^ neurogenesis defect. In turn, several mouse mutants defective for TZ proteins show microphthalmia (Garcia-Gonzalo et al., 2011; Sang et al., 2011; Yee et al., 2015). Tctn proteins are also required for Gli3 processing (Sang et al., 2011; Thomas et al., 2012; C. Wang, Li, Meng, & Wang, 2017) and the TZ protein Rpgrip1l controls the activity of the proteasome at the basal body responsible for proteolytic cleavage of Gli3 (Gerhardt et al., 2015). Taken together, these findings indicate that *Inpp5e* mutation might affect the ability of RGCs to switch to indirect neurogenesis through defects in cilia stability and/or the integrity of the ciliary transition zone.

### Implications for Joubert Syndrome

In humans, hypomorphic *INPP5E* mutations contribute to Joubert Syndrome (JS), a ciliopathy characterized by cerebellar malformations and concomitant ataxia and breathing abnormalities. In addition, a subset of JS patients exhibit cortical abnormalities including polymicrogyria, neuronal heterotopias and agenesis of the corpus callosum (Poretti, Huisman, Scheer, & Boltshauser, 2011). Strikingly, the *Inpp5e* mouse mutant also shows several of these abnormalities. In the caudal telencephalon, the otherwise lissencephalic cortex formed folds reminiscent of the polymicrogyria in JS patients. In addition, the mutant formed leptomeningeal heterotopias with 100% penetrance, but their number and location varied. Mutations in ciliary genes were previously associated with heterotopia formation in humans and mice (Magnani et al., 2015; Uzquiano et al., 2019). Mice carrying mutations in the *Eml1* gene encoding a microtubule-associated protein show subcortical heterotopias due to a mispositioning of radial glial cells and impaired primary cilia formation (Uzquiano et al., 2019). Finally, the corpus callosum is thinner but callosal axons normally project to the contralateral cerebral hemisphere in *Inpp5e* mutants. This phenotype is milder compared to that of other mouse mutants with altered cilia that show complete agenesis of the corpus callosum with callosal axons forming Probst bundles (Benadiba et al., 2012; Laclef et al., 2015; Putoux et al., 2019). Unlike these other ciliary mutants, the corticoseptal boundary which plays a crucial role in positioning guidepost cells that control midline crossing of callosal axons (Magnani et al., 2014) is not obviously affected in *Inpp5e*^Δ/Δ^ embryos. Instead, the thinner corpus callosum is likely to be the result of an expanded piriform cortex. Despite these mechanistic differences, however, re-introducing Gli3R into the cilia mutant background restores callosal development in both groups of mutants suggesting that cilia control two independent steps in corpus callosum formation by regulating Gli3 processing. Thus, the *Inpp5e*^Δ/Δ^ mutant recapitulates cortical abnormalities in JS patients and starts to help unravelling the pathomechanisms underlying these defects.

## MATERIAL & METHODS

### Mice

All experimental work was carried out in accordance with the UK Animals (Scientific Procedures) Act 1986 and UK Home Office guidelines. All protocols were reviewed and approved by the named veterinary surgeons of the College of Medicine and Veterinary Medicine, the University of Edinburgh, prior to the commencement of experimental work. *Inpp5e*^Δ/+^ and *Gli3*^Δ699/+^ mouse lines have been described previously (Böse et al., 2002; Jacoby et al., 2009). *Inpp5e*^Δ/+^ mice were interbred to generate *Inpp5e*^Δ/Δ^ embryos; exencephalic *Inpp5e*^Δ/Δ^ embryos were excluded from the analyses. Wild-type and *Inpp5e*^Δ/+^ litter mate embryos served as controls. *Inpp5e*^Δ/Δ^*Gli3*^Δ699/+^ and *Inpp5e*^Δ/Δ^*Gli3*^Δ699/Δ699^ embryos were obtained from inter-crosses of *Inpp5e*^Δ/+^;*Gli3*^Δ699/+^ mice using wild-type, *Inpp5e*^Δ/+^ and *Gli3*^Δ699/+^ embryos as controls. Embryonic (E) day 0.5 was assumed to start at midday of the day of vaginal plug discovery. Transgenic animals and embryos were genotyped as described (Böse et al., 2002; Jacoby et al., 2009). For each marker and each stage, 3-8 embryos were analysed.

For measuring cell cycle lengths, pregnant females were intraperitoneally injected with a single dose of IdU (10mg/ml) at E12.5, followed by an injection of BrdU (10mg/ml) 90 min later. Embryos were harvested 30 min after the second injection. For cell cycle exit analyses, BrdU was injected peritoneally into E11.5 pregnant females and embryos were harvested 24 hrs later. Rapamycin was dissolved in 100% EtOH and diluted in vehicle containing 5% PEG400, 5% Tween80 immediately before use. E11.5 pregnant females received a single intraperitoneal injection of Rapamycin at 1 mg/kg body weight or of vehicle and were killed 24 hrs later.

### Immunohistochemistry and in situ hybridisation

For immunohistochemistry, embryos were fixed overnight in 4% paraformaldehyde, incubated in 30% sucrose at +4°C for 24h, embedded in 30% sucrose/OCT mixture (1:1) and frozen on dry ice. Immunofluorescence staining was performed on 12 to 14 μm cryostat sections as described previously (Theil, 2005) with antibodies against Arl13b (Neuromab 75-287; 1:1500), rabbit anti-BrdU (1:50, Abcam #ab6326), mouse anti-BrdU/IdU (B44) (1:50, BD Biosciences #347580), rat anti-Ctip2 (1:1000, Abcam #18465), rabbit anti-pErk^T202/Y204^ (1:1000, Cell Signaling Technology #9101), rabbit anti-GFAP (1:1000, Dako #Z 0334), rat anti-L1, clone 324 (1:1000, Millipore #MAB5272), rabbit anti-Pax6 (1:400, Biolegend #901301), mouse anti-PCNA (1:500, Abcam #29), rabbit anti-Prox1 (1:1000, RELIA*Tech* #102-PA32). rabbit anti-pHH3 (1:100, Millipore #06-570), mouse anti-Satb2 (1:200, Abcam #51502), rabbit anti-Tbr1 (1:400, Abcam #31940), rabbit anti-Tbr2 (1:1000, Abcam #23345) and *γ*TUB (Sigma T6557; 1:2000). Primary antibodies for immunohistochemistry were detected with Alexa- or Cy2/3-conjugated fluorescent secondary antibodies. The Tbr1 signals were amplified using biotinylated secondary IgG antibody (swine anti-rabbit IgG) (1:400, BD Biosciences) followed by Alexa Fluor 488 or 568 Streptavidin (1:100, Invitrogen). For counter staining DAPI (1:2000, Life Technologies) was used. Prox1 and pErk^T202/Y204^ proteins were detected non-fluorescently using biotinylated goat anti-rabbit IgG (1: 400, BD Biosciences) followed by avidin-HRP and DAB detection (Vector labs) as described previously (Magnani et al., 2010).

In situ hybridisation on 12μm serial paraffin sections were performed as described previously (Theil, 2005) using antisense RNA probes for *Axin2* (Lustig et al., 2002), *Bmp4* (Jones, Lyons, & Hogan, 1991), *Dbx1* (Yun, Potter, & Rubenstein, 2001), *Dlx2* (Bulfone et al., 1993), *Emx1* (Simeone et al., 1992), *Gli3* (Hui, Slusarski, Platt, Holmgren, & Joyner, 1994), *Lhx2* (Liem, Tremml, & Jessell, 1997), *Msx1 (Hill et al., 1989)*, *Ngn2* (Gradwohl, Fode, & Guillemot, 1996), *Nrp2* (Galceran et al., 2000), *Pax6* (Walther & Gruss, 1991), *Scip1* (Frantz et al., 1994), *Wnt2b* (Grove, Tole, Limon, Yip, & Ragsdale, 1998).

### Western blot

Protein was extracted from the dorsal telencephalon of E12.5 (3 tissues pooled per sample) and E14.5 (single tissue per sample) wild-type and *Inpp5e*^Δ/Δ^ embryos (n=4 samples per genotype) as described previously (Magnani et al., 2010). For the detection of Gli3 10μg protein lysates were subjected to gel electrophoresis on a 3-8% NuPAGE® Tris-Acetate gel (Life Technologies), and protein was transferred to a Immobilon-FL membrane (Millipore), which was incubated with goat anti-h/m Gli3 (1:500, R&D Systems #AF3690) and mouse anti-β-Actin antibody (1:15000, Abcam #ab6276). After incubating with donkey anti-goat IgG IRDye680RD (1:15000, LI-COR Biosciences) and donkey anti-mouse IgG IRDye800CW secondary antibodies (1:15000, Life Technologies), signal was detected using LI-COR’s Odyssey Infrared Imaging System with Odyssey Software. Values for protein signal intensity were obtained using Image Studio Lite Version3.1. Gli3 repressor and activator protein levels were compared between wild-type and mutant tissue using a paired t-test.

For all other Western blot analyses 20μg protein lysates were loaded on 4–12% NuPAGE® Bis-Tris gels (Life Technologies) and later transferred to an Immobilon–FL membrane. Membranes were incubated with the following primary antibodies: rabbit anti-phospho-Akt^S473^ (1:1000, Cell Signaling Technologies CST #9271), rabbit anti-Akt (1:1000, CST #9272), rabbit anti-phospho-Erk^T202/Y204^ (1:1000, CST #9101), rabbit anti-Erk (L34F12) (CST # 4696), rabbit anti-ribosomal protein phospho-S6^S235/S236^ (1:1000, CST #2211), rabbit anti-ribosomal protein phospho-S6^S240/S244^ (1:1000, CST #2215) and mouse anti-ribosomal protein S6 (C-8) (1:1000, Santa Cruz Technologies #sc-74459). For the detection of phosphorylated proteins goat anti-rabbit IRDye 680RD (1:15,000, LI-COR Biosciences) and for total proteins goat anti-mouse or goat anti-rabbit IRDye 800CW (1:15,000, LI-COR Biosciences) were used as secondary antibodies. The signals were detected via the Odyssey Imaging System and further analysed using Image Studio Lite Version4.0. The ratios between phosphorylated and total protein were compated between wild-type and mutant tissue using a paired t-test.

### Scanning and transmission electron microscopy

TEM and SEM image acquisition were performed in the Cochin Imaging Facility and on the IBPS EM Facility, respectively. For scanning electron microscopy, embryos were dissected in 1.22x PBS (pH 7.4) and fixed overnight with 2% glutaraldehyde in 0.61x PBS (pH 7.4) at 4°C. Heads were then sectioned to separate the dorsal and ventral parts of the telencephalon, exposing their ventricular surfaces. Head samples were washed several times in 1.22x PBS and postfixed for 15 minutes in 1.22x PBS containing 1% OsO4. Fixed samples were washed several times in ultrapure water, dehydrated with a graded series of ethanol and prepared for scanning electron microscopy using the critical point procedure (CPD7501, Polaron). Their surfaces were coated with a 20 nm gold layer using a gold spattering device (Scancoat Six, Edwards). Samples were observed under a Cambridge S260 scanning electron microscope at 10 keV.

For transmission electron microscopy tissues were fixed for 1 hour with 3% glutaraldehyde, post-fixed in 1.22x PBS containing 1% OsO4, then dehydrated with a graded ethanol series. After 10 minutes in a 1:2 mixture of propane:epoxy resin, tissues were embedded in gelatin capsules with freshly prepared epoxy resin and polymerized at 60°C for 24 hours. Sections (80 nm) obtained using an ultramicrotome (Reichert Ultracut S) were stained with uranyl acetate and Reynold’s lead citrate and observed with a Philips CM10 transmission electron microscope.

### Statistical Analyses

Data were analysed using GraphPadPrism 6 software with n=3-8 embryos for all analyses. Power analysis of pilot experiments informed minimum samples size. Mann Whitney tests were performed for immunohistochemical analyses in general. Cortical thickness was analysed using a two way ANOVA followed by Sidak’s multiple comparisons test. Paired t-tests were performed for Western blots and a fisher’s exact test was used to analyse the quantification of normal and abnormal cilia. The mTORC1 and Gli3 rescue experiments were evaluated with one way ANOVAS followed by Tukey’s multiple comparisons test. A single asterisk indicates significance of p<0.05, two asterisks indicate significance of p<0.01 and three asterisks of p<0.005. Due to morphological changes blinding was not possible and scores were validated by a second independent observer.

## Supporting information

Summary of statistical analyses

## ACKNOWLEDGEMENTS

We are grateful to Drs Thomas Becker, John Mason, Pleasantine Mill, and David Price for critical comments on the manuscript, Dr Christos Gkogkas for invaluable advice on mTOR signaling, and Stéphane Schurmans for the *Inpp5e*^Δ/+^ mouse line. We also thank Dr Michaël Trichet (electron microscopy platform of the IBPS-Sorbonne Universités Paris 6) and Dr Alain Schmitt (electron microscopy platform of the Institut Cochin CNRS-UMR 8104) for their help with scanning and transmission electron microscopy analyses, respectively. This work was supported by a grant from the Biotechnology and Biological Sciences Research Council to TT (BB/P00122X/1) and NIH R01GM095941 to JFR.

