## Supplementary figures for "A transient role of primary cilia in controlling direct versus indirect neurogenesis in the developing cerebral cortex"

**Supplementary Figure 1: Formation of the telencephalic boundaries in *Inpp5e*$^{\Delta/\Delta}$ embryos.** (A-F) Formation of the corticoseptal boundary. Expression of the dorsal marker gene *Pax6* (A, B, D, E) and of the ventral marker gene *Dlx2* (C, F) remain restricted to the cortex and septum, respectively, with a sharp expression boundary between both tissues. (G-L) Formation of the pallial/subpallial boundary. While there is a sharp expression boundary between cortex and lateral ganglionic eminence (LGE) in wild-type embryos (G-I), scattered Pax6 and *Dlx2* expressing cells (arrows in J and K) are found in the mutant LGE and cortex, respectively, while the *Dbx1* expression domain characteristic of the ventral pallium (VP) is fuzzier (L). CGE: caudal ganglionic eminence; ctx: cortex; MGE: medial ganglionic eminence; sep: septum; th: thalamus. Scale bars: 200μm.

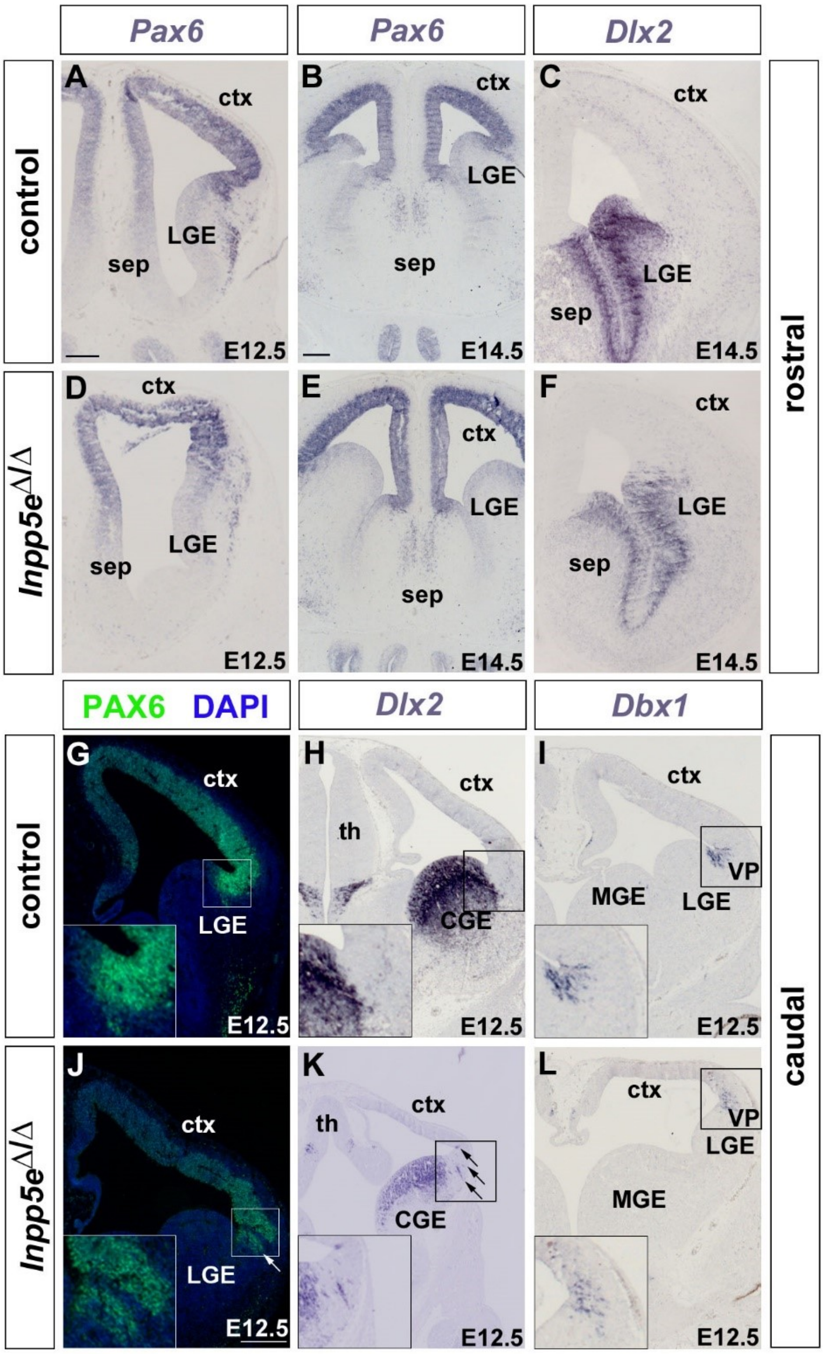

**Supplementary Figure 2: Wnt/β-catenin and Bmp signalling in the dorsomedial telencephalon of E12.5 *Inpp5e*$^{\Delta/\Delta}$ embryos.** (A, B, E, F) Reduced Wnt/β-catenin signalling in *Inpp5e*$^{\Delta/\Delta}$ embryos. (A, E) *Wnt2b* expression is confined to the cortical hem (h) while there are only few scattered Wnt2b expressing cells in the mutant. (B, F) Graded expression of the Wnt target gene *Axin2* in the dorsal midline is reduced in mutant embryos. (C, D, G, H) Roof plate (rp) expression of *Bmp4* and its target gene *Msx1* are reduced in *Inpp5e*$^{\Delta/\Delta}$ embryos. Scale bar: 100μm.

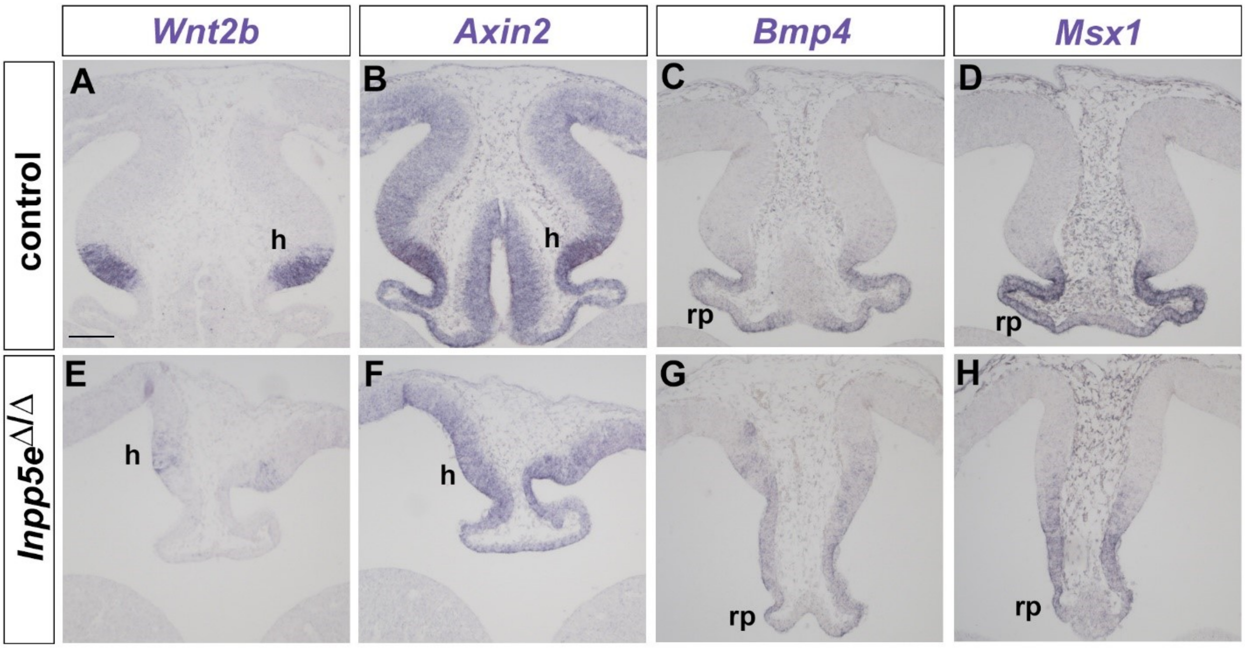

**Supplementary Figure 3: Expression of cortical progenitor markers in *Inpp5e*$^{\Delta/\Delta}$ embryos.** (A-J) Dorsal marker gene expression at patterning stages (E12.5). *Emx1*, *Lhx2*, *Pax6* and *Ngn2* are still expressed in the developing neocortex of *Inpp5e*$^{\Delta/\Delta}$ embryos though the *Lhx2* medial to lateral (B, C, G, H) and the *Pax6*/*Ngn2* lateral to medial (D, E, I, J) expression gradients are flatter. (K-T) Neocortical progenitors express *Lhx2*, *Pax6* and *Ngn2* in E14.5 *Inpp5e*$^{\Delta/\Delta}$ embryos. Note the folding of the neocortex at caudal levels (arrows in P, R and T). CGE: caudal ganglionic eminence; ctx: cortex; MGE: medial ganglionic eminence; LGE: lateral ganglionic eminence; pt: prethalamus; sep: septum; th: thalamus. Scale bars: 200μm.

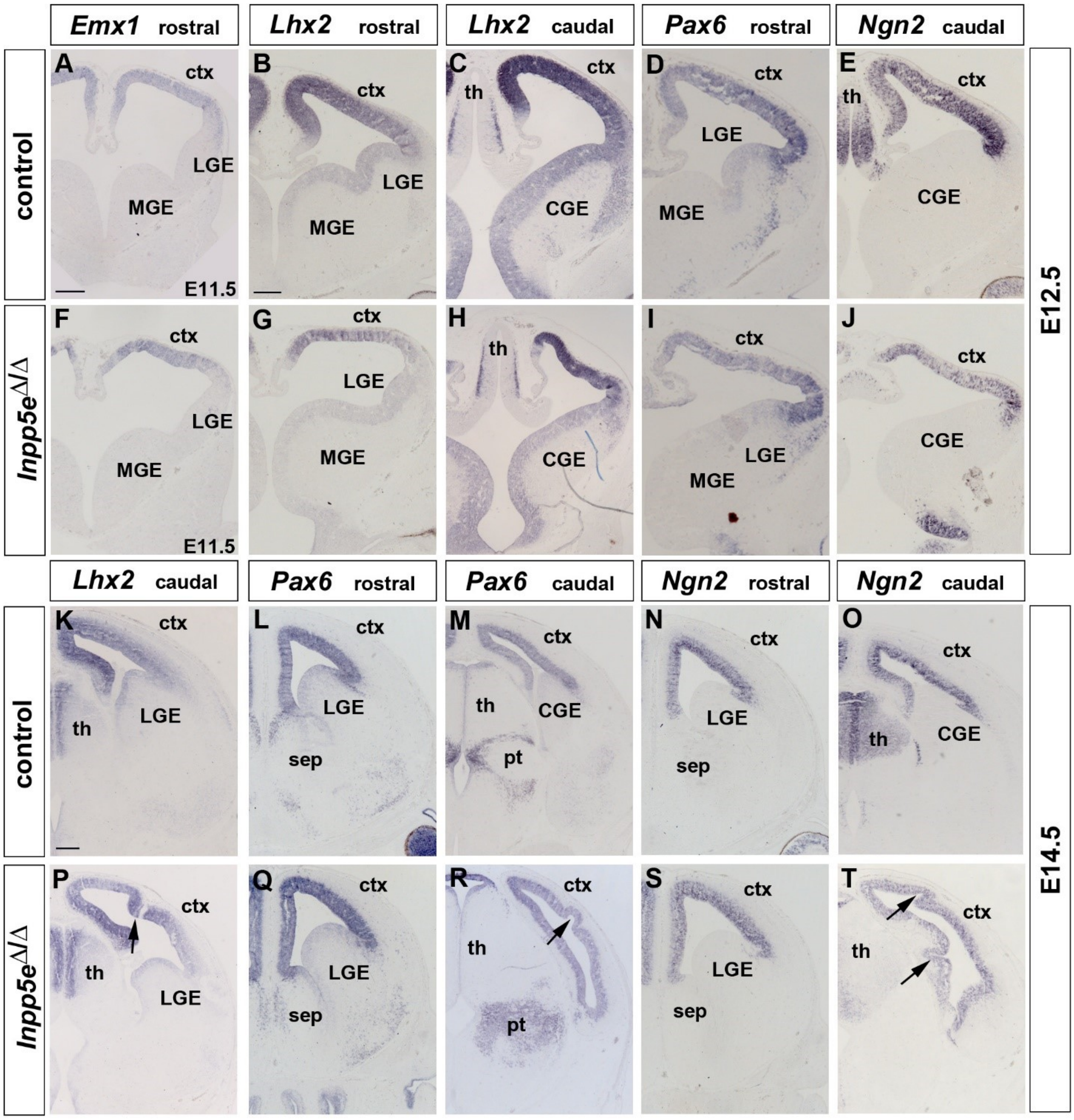

**Supplementary Figure 4: Proportion of mitotic progenitors in *Inpp5e*$^{\Delta/\Delta}$ embryos.** (A-H) Proportions of mitotic progenitors in E12.5 control (A, E) and *Inpp5e*$^{\Delta/\Delta}$ embryos (B, F) as revealed by pHH3 (mitotic cells) and PCNA (all progenitor cells) double immunofluorescence. Note the reduction in mitotic basal progenitors in the *Inpp5e*$^{\Delta/\Delta}$ medial neocortex (A, B, D). (I-P) The proportions of apical and basal progenitors is not significantly different in E14.5 control and *Inpp5e*$^{\Delta/\Delta}$ embryos. In all panels, radial glia cells divide at the ventricular surface whereas mitotic basal progenitors locate in abventricular positions. All statistical data are presented as means ± 95% confidence intervals (CI); Mann Whitney tests; n = 4; * $p < 0.05$. Scale bar: 50μm.

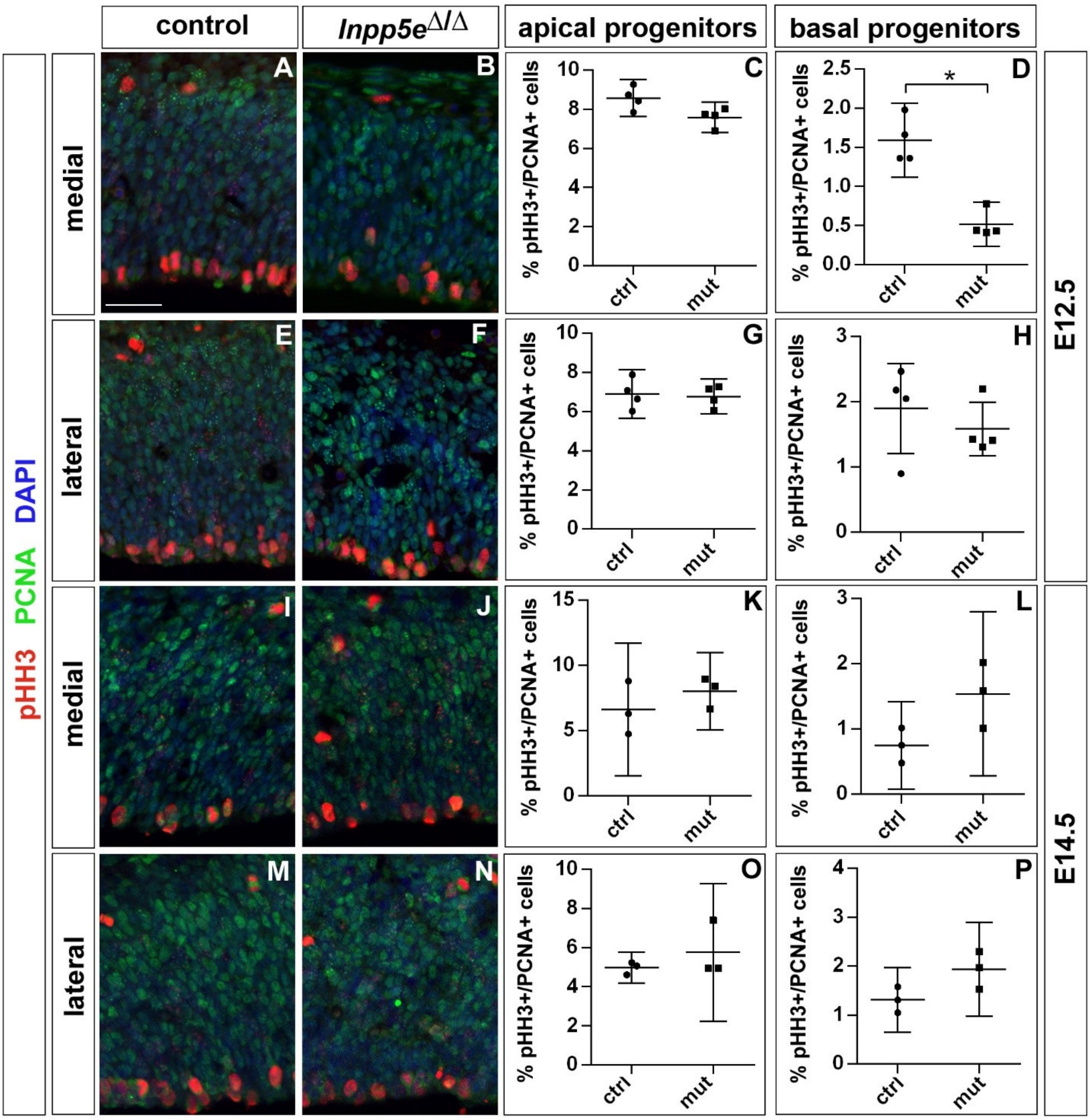

**Supplementary Figure 5: Cell cycle of cortical progenitors in E12.5 *Inpp5e*$^{\Delta/\Delta}$ embryos.** (A) Schematic illustrating the BrdU/IdU double labelling strategy to measure S phase ($T_S$) and total cell cycle length ($T_C$). 90 minutes after an initial IdU administration, pregnant females received an intraperitoneal BrdU injection. Embryos are harvested 30 minutes later. (B, C) Double immunofluorescence to detect IdU+ and BrdU+ progenitors. (D) Quantification showing no significant change in $T_S$ and $T_C$. Statistical data are presented as means ± 95% confidence intervals (CI); Mann Whitney tests; n = 4; * $p < 0.05$. Scale bar: 50μm.

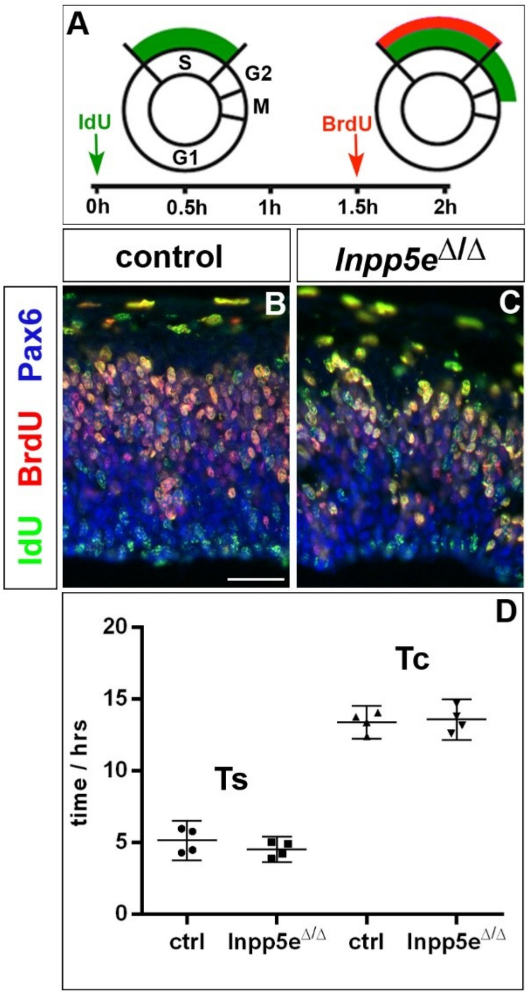

**Supplementary Figure 6: Whole mount preparations of E18.5 brains.** (A) Control. (B) *Inpp5e*$^{\Delta/\Delta}$ brain. Note the absence of obvious protrusions of the olfactory bulbs (ob) in the mutant. Ctx: cortex. Scale bar: 1 mm.

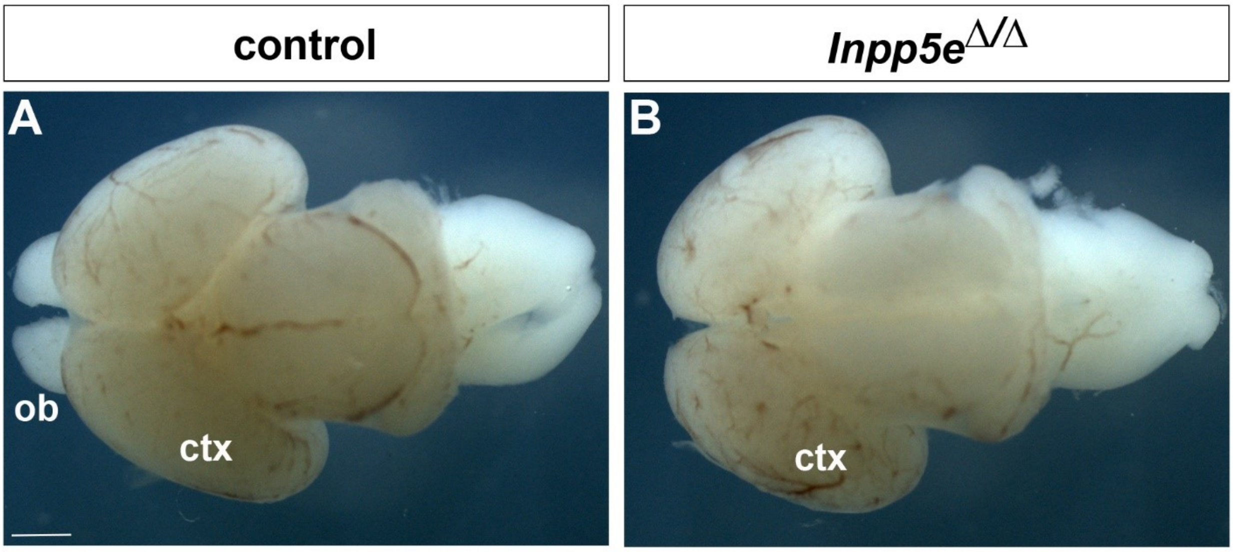

**Supplementary Figure 7: Forebrain malformations in E18.5 *Inpp5e*$^{\Delta/\Delta}$ embryos.** (A-F) Coronal sections through the forebrain of E18.5 control (A-C) and *Inpp5e*$^{\Delta/\Delta}$ (D-F) embryos. The asterisk in (D) demarcates a heterotopia in the *Inpp5e* mutant. Note that the mutant lateral neocortex is thinner at intermediate and caudal levels (E, F). The lines in (A) indicate where cortical thickness was measured at medial (m) and lateral (l) levels. (G, H) Quantification of cortical thickness. CC: corpus callosum; ctx: cortex; hip: hippocampus; sep: septum; str: striatum; th: thalamus. Scale bar: 500μm. Statistical data are presented as means ± 95% confidence intervals (CI); Two way ANOVA followed by Sidak multiple comparisons test; n = 4; * $p < 0.05$; ** $p < 0.005$; *** $p < 0.001$.

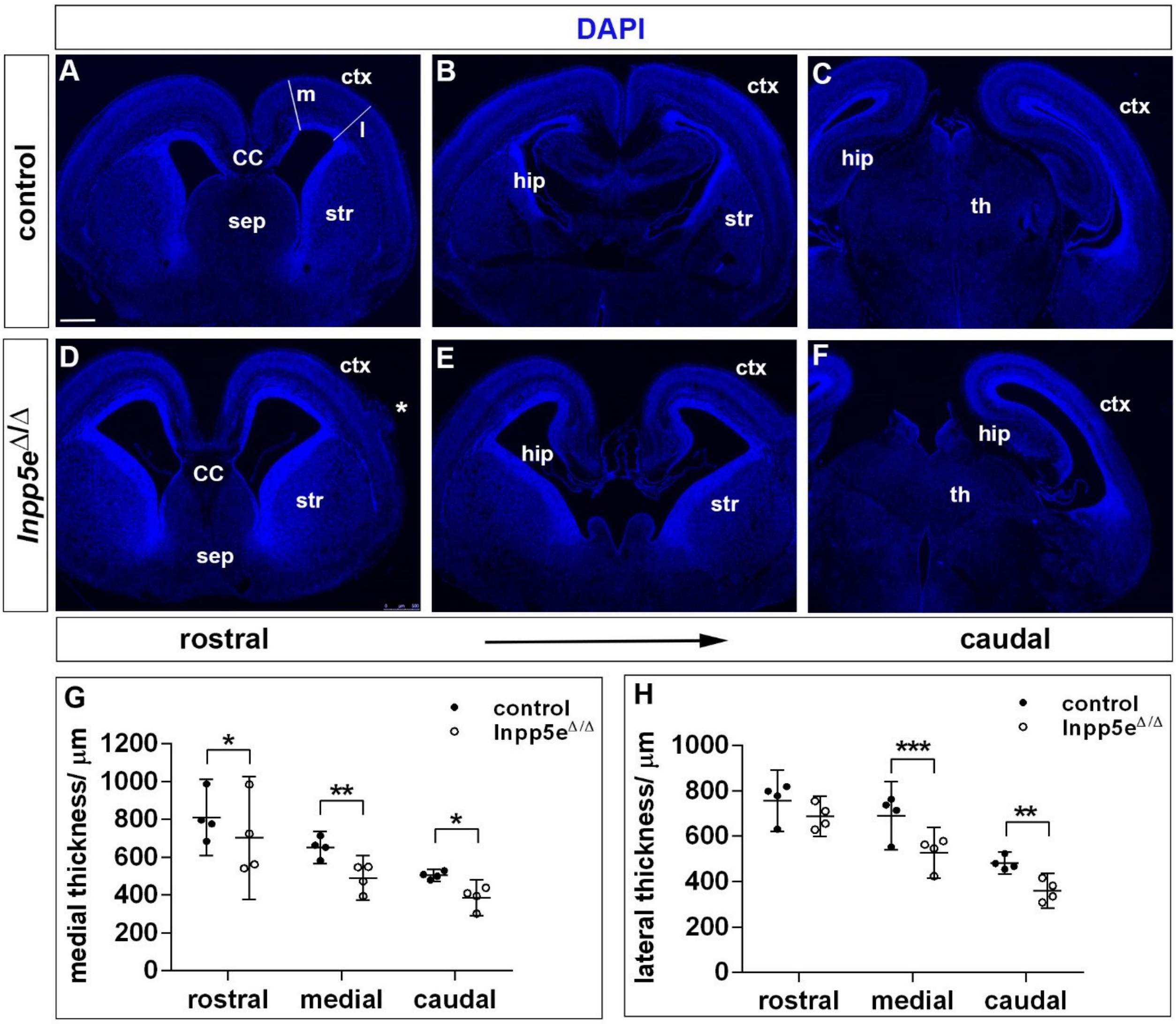

**Supplementary Figure 8: Hippocampus formation in *Inpp5e*$^{\Delta/\Delta}$ embryos.** (A-F) Hippocampal marker gene expression in E18.5 control (A-C) and *Inpp5e*$^{\Delta/\Delta}$ embryos (D-F). Expression of *Nrp2* labels the whole hippocampal formation (A), while *Scip1* is expressed in CA1 and in the neocortex (B). Prox1 expression is confined to the dentate gyrus (DG) (C). In *Inpp5e*$^{\Delta/\Delta}$ embryos, these hippocampal markers are expressed but their expression domains are severely reduced or disorganized (D-F).

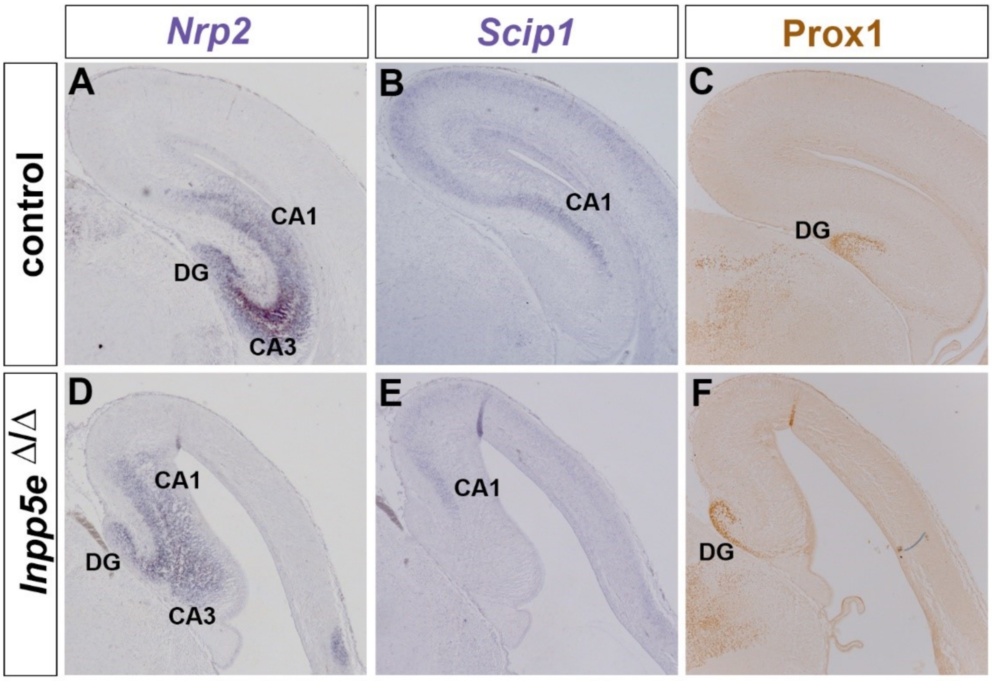


**Supplementary Figure 9: Formation of the corpus callosum in E18.5 *Inpp5e*$^{\Delta/\Delta}$ embryos.** (A, B) Coronal section through the telencephalon stained with L1 and GFAP to reveal the corpus callosum (CC) and glial cells, respectively. The corpus callosum is smaller in *Inpp5e*$^{\Delta/\Delta}$ embryos while the glial wedge (GW), the induseum griseum glia (IGG) and the midline zipper glia (MZG) occupy their correct position surrounding the corpus callosum. (C) Quantification of corpus callosum thickness. Statistical data are presented as means ± 95% confidence intervals (CI); Mann Whitney tests; n = 4; * $p < 0.05$. Scale bar: 250µm.

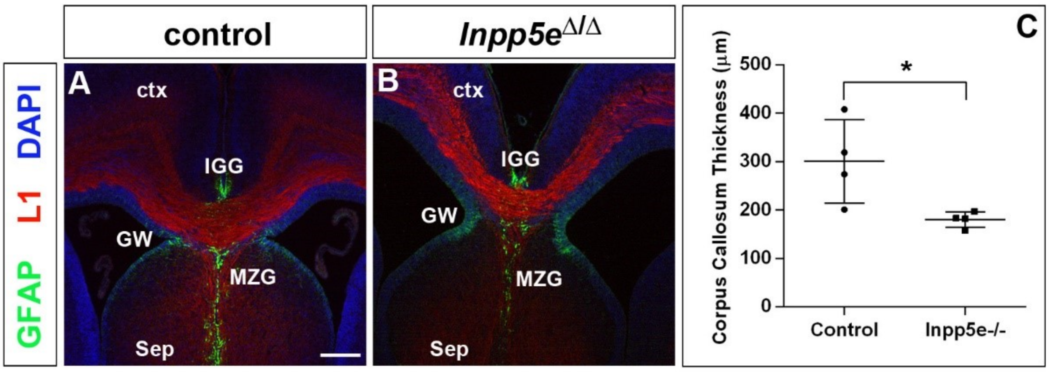

**Supplementary Figure 10: Erk signalling in *Inpp5e*$^{□/□}$ embryos.** (A, E) Western blot analysis on dorsal telencephalic tissue from E12.5 (A) and E14.5 (E) embryos using anti-phosphoErk$^{T202/Y204}$ and total-Erk antibodies. (B, F) Quantification of the Western blots. A paired t-test was used to evaluate relative pErk/Erk expression levels in four control/*Inpp5e*$^{□/□}$ embryo pairs derived from four different litters; statistical data are presented as means ± 95% confidence intervals (CI). (C, D, G, H) Immunohistochemistry for phosphoErk$^{T202/Y204}$ on coronal sections of the telencephalon of E12.5 (C, D) and E14.5 (G, H) embryos. ctx: cortex; MGE: medial ganglionic eminence; LGE: lateral ganglionic eminence; th: thalamus Scale bars: 100 μm.

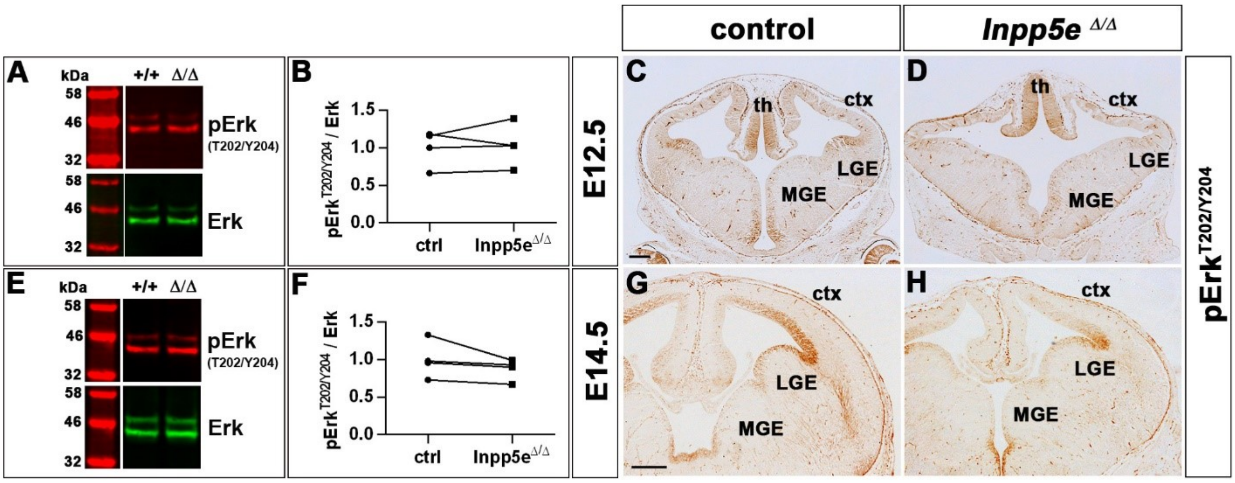

**Supplementary Figure 11: Akt and mTOR signalling in *Inpp5e*$^{\Delta/\Delta}$ embryos.** (A, E) Western blot analyses on dorsal telencephalic tissue from E12.5 (A, C, E) and E14.5 (B, D, F) embryos with the indicated antibodies. (A, B) Increased ratio of pAkt$^{S473}$/total Akt in E12.5 *Inpp5e*$^{\Delta/\Delta}$ embryos but not at E14.5. (C, D) Phosphorylation of S6 at residues S235/S236 is increased in the dorsal telencephalon of E12.5 and E14.5 *Inpp5e*$^{\Delta/\Delta}$ embryos. (E, F) E12.5 *Inpp5e*$^{\Delta/\Delta}$ embryos show an increased phosphorylation of S6 at residues S240/S244. (G-P) *Inpp5e*$^{\Delta/\Delta}$ embryos treated with rapamycin show an augmented proportion of Tbr1+ neurons (G-J, O) and a reduced proportion of Tbr2+ basal progenitors (K-N, P). For quantification of the Western blots, paired t-test were used to evaluate expression levels of the phosphorylated versus unphosphorylated proteins in four control/*Inpp5e*$^{\Delta/\Delta}$ embryo pairs derived from four different litters. Immunofluorescence stainings were analysed by Mann Whitney tests. Statistical data are presented as means ± 95% confidence intervals (CI); n = 4; * $p < 0.05$; *** $p < 0.005$.

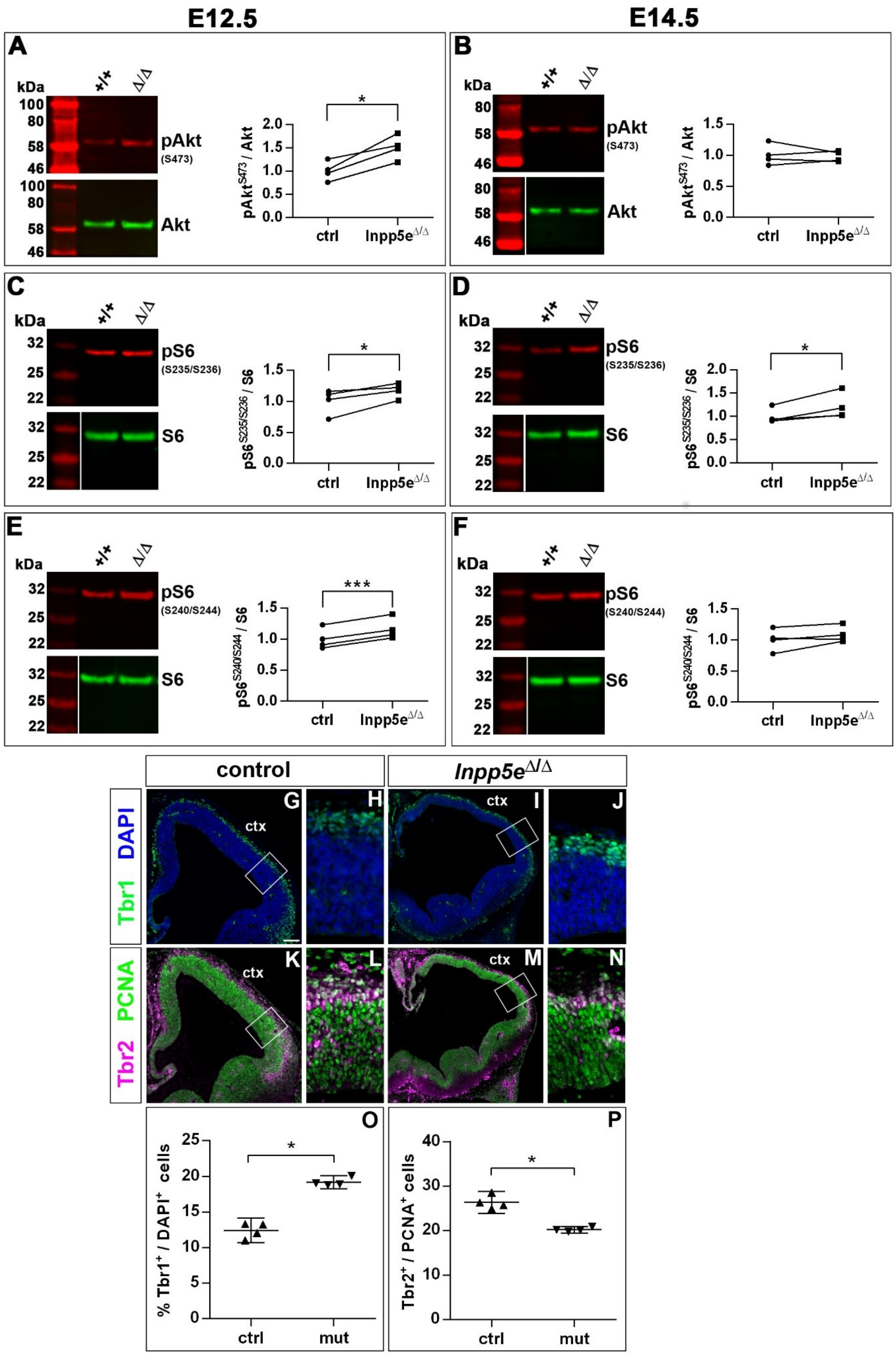

E12.5
E14.5
A
kDa
+/+
Δ/Δ
100
80
58
46
pAkt
(S473)
Akt
*
2.0
1.5
1.0
0.5
0.0
pAkt S473 / Akt
ctrl
Inpp5e Δ/Δ
B
kDa
80
58
46
1.5
C
kDa
32
25
22
pS6
(S235/S236)
S6
pS6 S235/S236 / S6
D
E
***
pS6
(S240/S244)
pS6 S240/S244 / S6
F
control
Inpp5e Δ/Δ
DAPI
Tbr1
PCNA
Tbr2
ctx
G
H
I
J
K
L
M
N
O
P
25
20
15
10
5
0
% Tbr1+ / DAPI+ cells
ctrl
mut
40
30
20
10
0
Tbr2+ / PCNA+ cells

**Supplementary Figure 12: *Gli3* mRNA expression in Inpp5e mutants.** (A, B) *Gli3* in situ hybridization showing *Gli3* mRNA expression in the cortex (ctx) and lateral ganglionic eminence (LGE) of control (A) and *Inpp5e*$^{\Delta/\Delta}$ embryos (B).

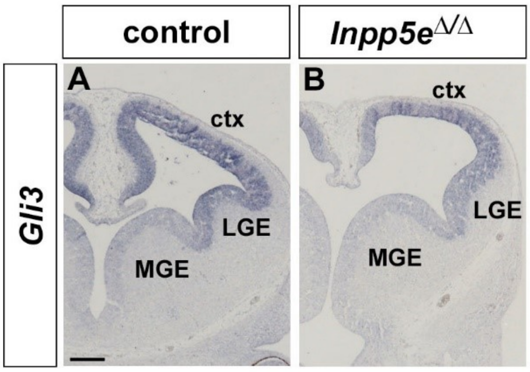

**Supplementary Figure 13: Rescue of eye development in *Inpp5e*$^{\Delta/\Delta}$;*Gli3*$^{\Delta699/+}$ embryos.** (A-C) Side views of the heads of E12.5 embryos with the indicated genotype. *Inpp5e*$^{\Delta/\Delta}$ embryos lack the eye completely or only form a small remnant whereas eye formation is not affected in *Inpp5e*$^{\Delta/\Delta}$;*Gli3*$^{\Delta699/+}$ embryos. Scale bar: 1mm

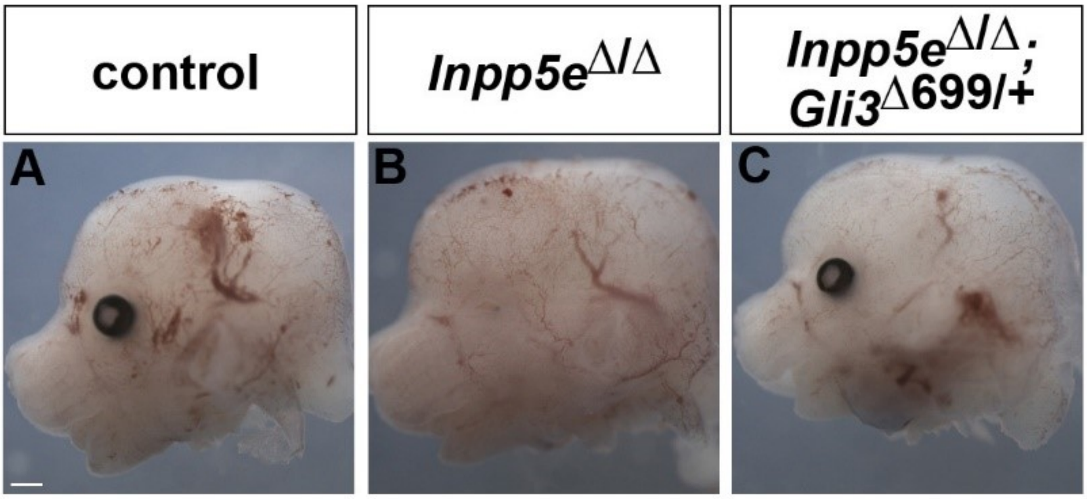


**Supplementary Figure 14: Rescue of corpus callosum formation in *Inpp5e*$^{\Delta/\Delta}$;*Gli3*$^{\Delta699/+}$ embryos.** (A-B) Coronal section through the telencephalon stained with L1 and GFAP to reveal the corpus callosum (CC) and glial cells, respectively. There is no significant difference in the size of the corpus callosum between control and *Inpp5e*$^{\Delta/\Delta}$;*Gli3*$^{\Delta699/+}$ embryos; the glial wedge (GW), the induseum griseum glia (IGG) and the midline zipper glia (MZG) are formed in their correct position. (C) Quantification of corpus callosum thickness. Statistical data are presented as means ± 95% confidence intervals (CI); Mann Whitney tests; n = 4; Scale bar: 250µm.

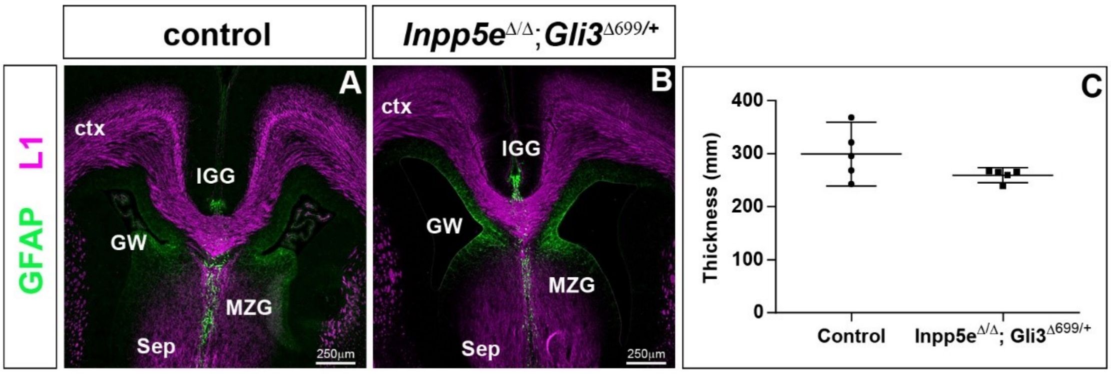
